# Design, Synthesis, Biological Evaluation and Molecular Docking of New Acid-Functionalized Carbazole Derivatives as Potential Antibiotic Agents

**DOI:** 10.1101/2025.09.11.675744

**Authors:** Aniruddhya Mukherjee, Ananya Das Mahapatra, Khushhali Menaria Pandey, Aditya Maity, Suchandra Chakraborty

## Abstract

The emergence of antibiotic resistance necessitates the discovery of new scaffolds with improved efficacy and drug-like properties. In this work, a new series of acid-functionalized carbazole derivatives was designed, synthesized, and comprehensively evaluated for their antibacterial potential. The incorporation of acidic functionalities into the carbazole framework enhanced physicochemical properties and biological interactions, yielding distinct strain-specific antibacterial activities. Minimum inhibitory concentration (MIC) studies demonstrated that 3-methyl-1,4-dioxo-4,9-dihydro-1*H*-carbazole-6-carboxylic acid (**2**) exhibited broad-spectrum potency, particularly against *S. aureus* and *E. coli*, while 6-methyl-9*H*-carbazole-3-carboxylic acid (**3**) showed selectivity against *B. cereus* and (*E*)-3-methyl-1-(2-tosylhydrazono)-2,3,4,9-tetrahydro-1*H*-carbazole-6-carboxylic acid (**1**) displayed strong activity toward *S. typhimurium*. Molecular docking studies revealed favourable binding affinities of all derivatives toward bacterial dihydrofolate reductase (DHFR), with compound **1** showing the highest docking scores. Molecular dynamics simulations further confirmed the broad conformational adaptability of compound **1**, the target-specific stability of compound **3**, and protein-dependent binding behaviour of compound **2**. Complementary ADMET predictions indicated that all compounds adhered to Lipinski’s rules, with compound **3** displaying the most favourable pharmacokinetic profile, including high oral bioavailability and low toxicity risk. Together, these experimental and computational findings establish acid-functionalized carbazole scaffolds as promising antibacterial candidates.

## Introduction

The rapid and widespread emergence of drug-resistant bacterial strains has become one of the most pressing global health challenges of the 21st century. Antibacterial resistance is now well recognized as a major contributor to increased morbidity, mortality, and healthcare costs, significantly compromising the effectiveness of existing antibiotics. In particular, hospital-acquired infections caused by Gram-positive and Gram-negative pathogens—such as *Staphylococcus aureus* and *Escherichia coli*—are becoming increasingly difficult to treat due to their resistance to multiple drug classes [1–3]. Methicillin-resistant *Staphylococcus aureus* (MRSA) and multidrug-resistant (MDR) *E. coli* exemplify the severity of this crisis, making routine infections potentially life-threatening and undermining clinical outcomes [4–7]. The World Health Organization has declared antimicrobial resistance a top global public health threat, with current estimates suggesting that multidrug-resistant bacteria are responsible for over 400 million deaths annually [8,9]. These alarming statistics underscore the urgent need for novel antimicrobial agents with improved efficacy and reduced susceptibility to resistance.

There has been significant interest among chemists and biologists in carbazole alkaloids owing to their unique structural characteristics and promising pharmacological potential [10–13], after the initial isolation of Murrayanine, 3-formyl-1-methoxycarbazole, a carbazole alkaloid having antibiotic properties from *Murraya koenigii* Spreng [14–16]. Carbazole derivatives characterized by a *N*-containing rigid aromatic heterocyclic core with favourable electronic charge transfer properties [17], have attracted significant attention in medicinal chemistry due to their diverse biological activities, including antibacterial [18,19], antifungal [20,21], antitumor [22], and anti-inflammatory properties. Among these, carbazole-1,4-quinones have demonstrated the most potent anti-tubercular (anti-TB) activity [23,24].

The field of carbazole chemistry has witnessed significant advancements following the isolation of carbazomycins by Nakamura and co-workers from *Streptoverticillium ehimense* H 1051-MY10 (Figure 1). Among them, Carbazomycin A (1) and Carbazomycin B (2) have demonstrated potent antibacterial and antifungal activities, notably inhibiting the growth of phytopathogenic fungi and exhibiting both antibacterial and antiyeast properties [25]. In addition, Carbazomycin B (2) and Carbazomycin C (3) have been identified as inhibitors of 5-lipoxygenase, an enzyme linked to inflammatory processes [26]. Carbazomycin G (7) has shown antifungal activity specifically against *Trichophyton* species [27]. The biological activities of these natural products are largely influenced by the nature and position of functional groups on the carbazole scaffold. For instance, the naturally occurring carbazoloquinone alkaloid Murrayaquinone A, which possesses a quinone moiety and a methyl substituent at the C-3 position, has exhibited cardiotonic activity in guinea pig papillary muscle assays [28]. Carbazomycins G and H, featuring a methyl group at the C-2 position and a distinct quinol functionality, also contribute to the structural diversity relevant to bioactivity. Furthermore, it has been reported that introducing functional groups such as –CH□OH, –CHO, – COOH, and –COOCH□ enhances the selective bioactivity of 3-methylcarbazole derivatives [28, 29]. The presence of electron-withdrawing groups on the aromatic ring is known to improve the polarity, solubility, and stability of the carbazole nucleus, thereby enhancing its pharmacological potential [29].

**Figure 1.**
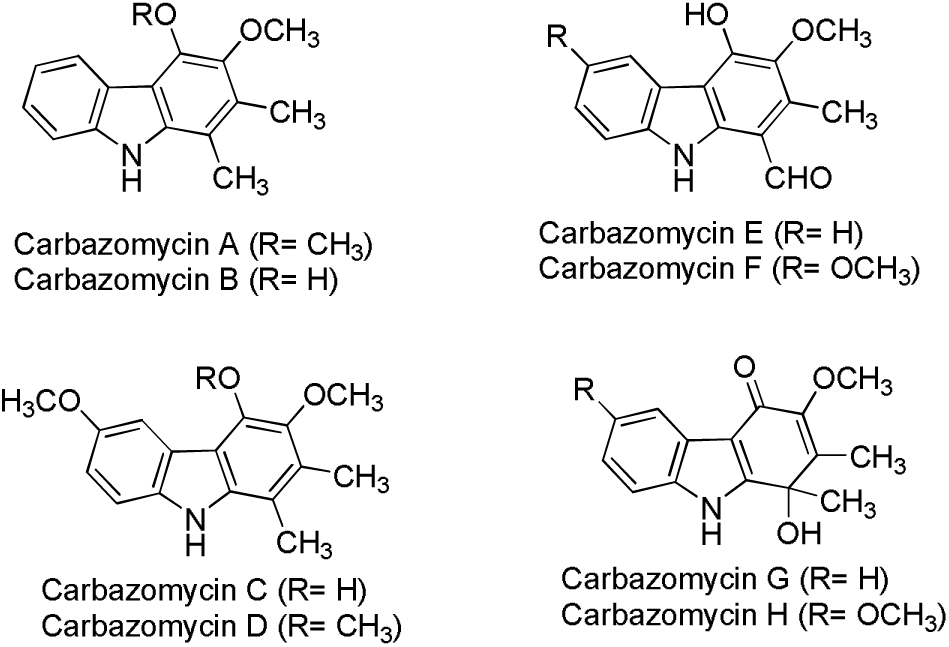
Naturally occurring Carbazomycins

**Figure 2.**
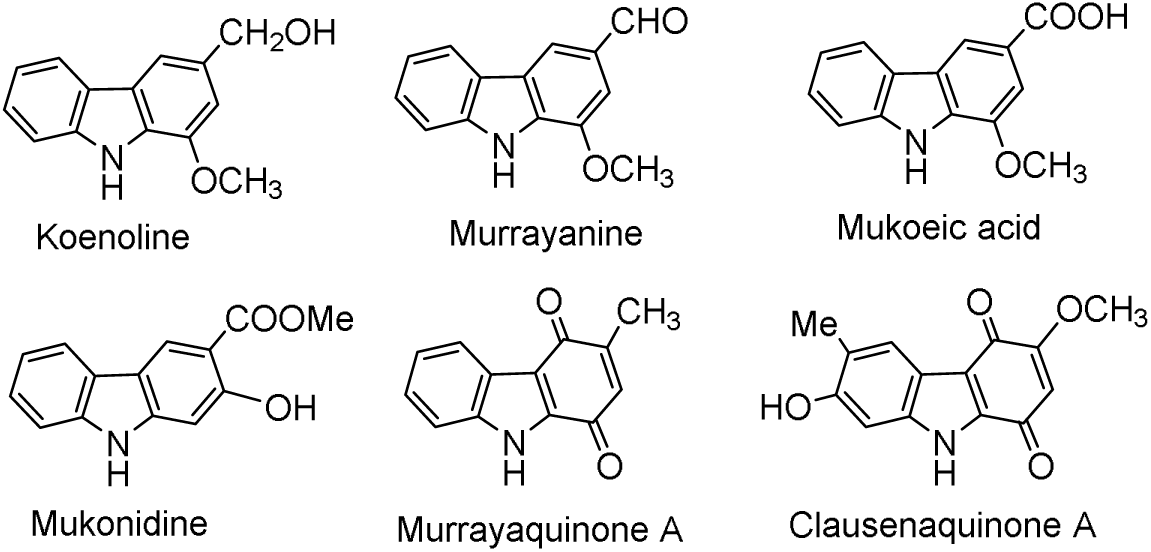
Natural carbazoles with various functional groups

Since the development of carbapenem antibiotics such as imipenem [30], which served as the parent scaffold for all subsequent carbapenem derivatives, the functionalization of drug molecules with acidic groups has emerged as a valuable strategy in the drug discovery process. Several clinically important antibiotics contain carboxylic acid functionalities critical to their pharmacological activity and pharmacokinetics, e.g. β-lactam antibiotics such as penicillins and cephalosporins possess carboxyl groups essential for their interaction with bacterial transpeptidases, thereby inhibiting cell wall synthesis [31]. Fluoroquinolone antibiotics such as ciprofloxacin and levofloxacin also contain carboxylic acid moieties that facilitate binding to bacterial DNA gyrase and topoisomerase IV [32]. Numerous studies have demonstrated that the incorporation of carboxylic acid or related acidic functionalities into bioactive compounds can significantly enhance their pharmacological profiles and therapeutic efficacy. The presence of an acid group increases aqueous solubility, thereby facilitating cellular absorption. Additionally, its electron-withdrawing nature modulates the basicity of the molecule, often enhancing chemical reactivity and improving interactions with biological targets. The influence of pKa on pharmacokinetic parameters and binding affinity is well-established, as it directly affects ionization states and molecular interactions within physiological environments. Acidic groups also contribute to increased hydrophilicity, which is advantageous for drug transport and bioavailability. In the context of rational drug design, achieving an optimal balance between polarity and hydrophilicity is essential for in vivo efficacy.

Designing novel compounds that are structurally distinct yet closely mimic the key features of known bioactive molecules is a cornerstone of medicinal chemistry. In our study so far, we have reported the high antibacterial efficacy of the various synthesised naturally occurring carbazole derivatives [33,34] along with some chloro and fluoro carbazole and carbazoloquinones [35, 36]. The carbazole scaffold, with its rigid aromatic system and modifiable framework, offers remarkable synthetic flexibility and potential for tuning physicochemical properties. In particular, acid-functionalized carbazole derivatives represent a promising class of therapeutic candidates. The introduction of acidic moieties can enhance solubility, improve pharmacokinetic behaviour, and promote target engagement through hydrogen bonding and ionic interactions. Despite their potential, no systematic study has yet been reported on the synthesis, and biological evaluation of acid-functionalized carbazole derivatives as antibacterial agents.

**Figure 3.**
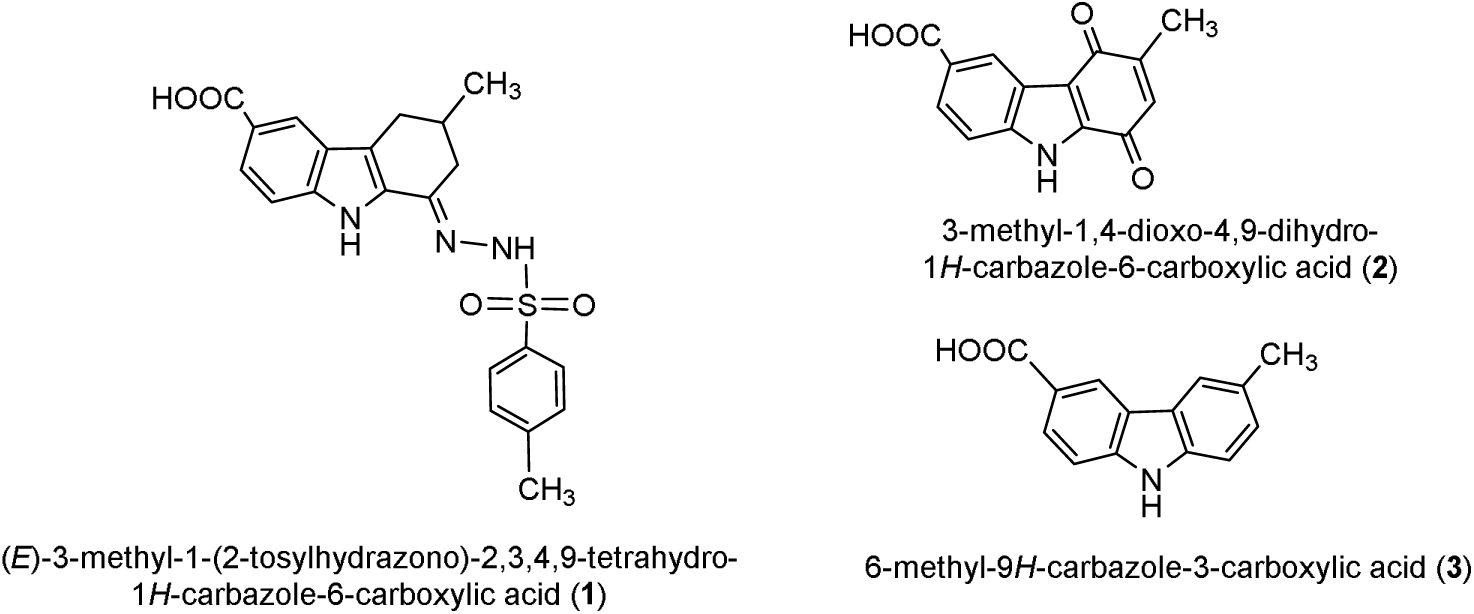
Design of acid functionalised carbazole derivatives

In this context, we aimed to explore this promising chemical space by designing and synthesising novel acid-functionalized carbazole-based derivatives, specifically 6-methyl-9*H*-carbazole-3-carboxylic acid analogues. The targeted compounds included (*E*)-3-methyl-1-(2-tosylhydrazono)- 2,3,4,9-tetrahydro-1*H*-carbazole-6-carboxylic acid (**1**), 3-methyl-1,4-dioxo-4,9-dihydro-1*H*-carbazole-6-carboxylic acid (**2**) and 6-methyl-9*H*-carbazole-3-carboxylic acid (**3**). The design was guided by considerations of structure–activity relationships (SAR), functional group-assisted reactivity, the oxidizing nature of tosyl and quinone functionalities, and the influence of electronegative groups on molecular polarity. The synthesized compounds were evaluated for their in vitro antibacterial activity against both Gram-positive and Gram-negative strains like *Bacillus cereus* (MTCC 430), *Staphylococcus aureus* (MTCC 87), *Escherichia coli* (MTCC 46) and *Salmonella typhimurium* (MTCC 733). Furthermore, molecular docking studies were conducted to provide insight into the binding interactions of these compounds with bacterial target proteins, aiming to rationalize their observed biological profiles. The integration of synthetic chemistry, biological evaluation, and in silico modelling in this study may contribute to the development of promising lead structures in the fight against antimicrobial resistance.

## Result and Discussion

### Chemistry

As a continuation of our synthetic efforts on carbazole alkaloids, we focused on further functionalizing the readily accessible 2,3,4,9-tetrahydro-1*H*-carbazol-1-one scaffold by introducing a carboxylic acid (-COOH) group onto the aromatic ring to enhance the biological activity of the system. In this context, we designed the synthesis of 3-methyl-1-oxo-2,3,4,9-tetrahydro-1*H*-carbazole-6-carboxylic acid (**4**), starting from commercially available 2-methylcyclohexanone (**5**) and *p*-aminobenzoic acid. The synthetic route commenced with a Claisen condensation [37] of compound **5** with ethyl formate in the presence of metallic sodium and a catalytic amount of ethanol in anhydrous ether, yielding 3-methyl-2-oxocyclohexanecarbaldehyde (**6**). Subsequent condensation of compound **6** with the diazonium salt derived from *p*-aminobenzoic acid (**7**), under Japp–Klingemann conditions [38], afforded the hydrazone intermediate (*E*)-4-(2-(4-methyl-2-oxocyclohexylidene)hydrazinyl)benzoic acid (**8**). Finally, Fischer indole cyclization [38] of compound **8** was carried out by refluxing in glacial acetic acid in the presence of concentrated hydrochloric acid for 3 minutes, resulting in the formation of the target compound **4** (Scheme 1), which was confirmed by the ¹H NMR spectrum,□displayed characteristic signals at δ 11.85 for –COOH and methylene protons as multiplet in the region 2.46–2.60 ppm. Again, ¹³C NMR spectrum□showed key resonances at δ 191.16 and 168.54 ppm for keto (C=O) and –COOH groups. This synthetic strategy highlights the successful incorporation of a carboxylic acid group into the carbazole core, setting the stage for further extension.

**Scheme 1.**
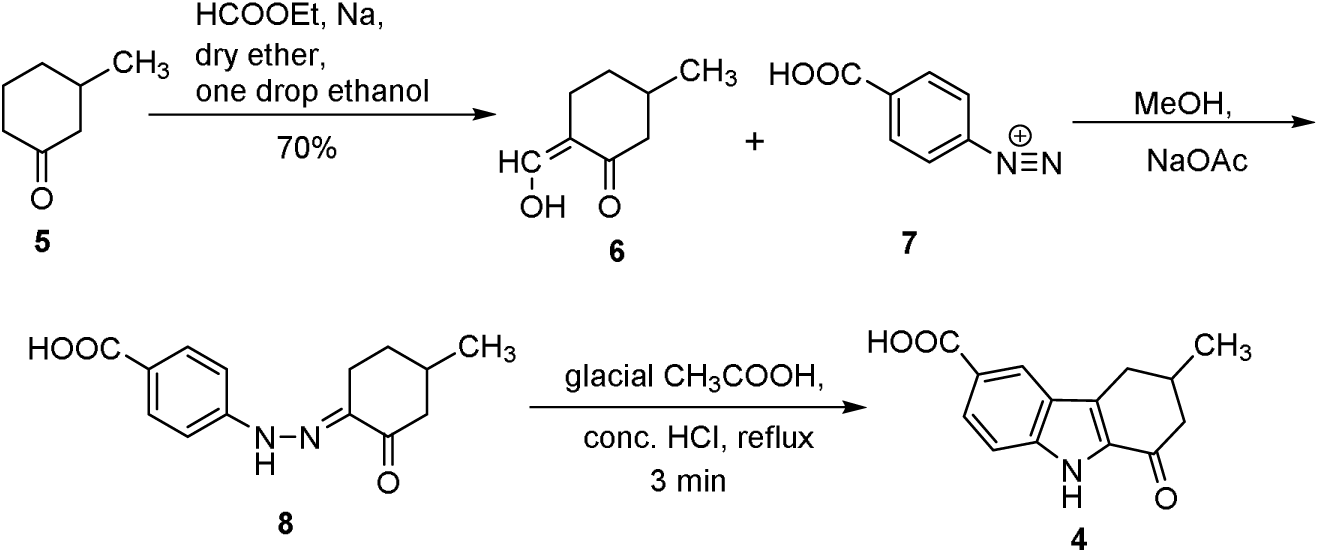
Synthesis of 3-methyl-1-oxo-2,3,4,9-tetrahydro-1*H*-carbazole-6-carboxylic acid (**4**)

The target compounds **1**, **2**, and **3** were efficiently synthesized from the key intermediate, 3-methyl-1-oxo-2,3,4,9-tetrahydro-1*H*-carbazole-6-carboxylic acid (**4**). Initially, (*E*)-3-methyl-1-(2-tosylhydrazono)-2,3,4,9-tetrahydro-1*H*-carbazole-6-carboxylic acid (**1**) was obtained in excellent yield by refluxing compound **4** with *p*-toluenesulfonylhydrazide in methanol for 5 hours (Scheme 2) [39]. The resulting product crystallized directly from the reaction mixture upon standing overnight under cold conditions.

**Scheme 2.**
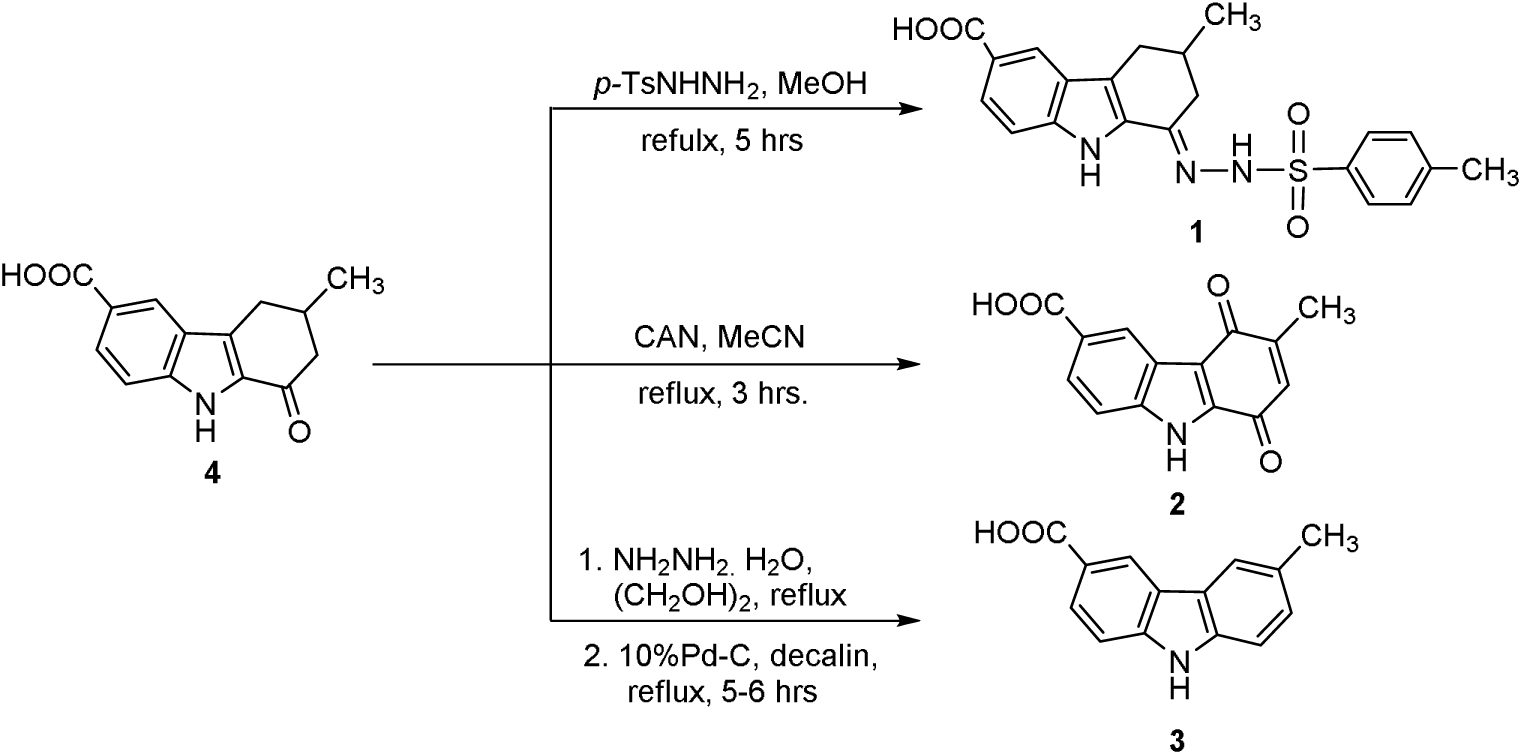
Synthesis of (*E*)-3-methyl-1-(2-tosylhydrazono)-2,3,4,9-tetrahydro-1*H*-carbazole-6-carboxylic acid (**1**), 3-methyl-1,4-dioxo-4,9-dihydro-1*H*-carbazole-6-carboxylic acid (**2**) and 6-methyl-9*H*-carbazole-3-carboxylic acid (**3**).

Subsequently, the 6-carboxylic acid-substituted carbazoloquinone derivative (**2**) was synthesized by oxidative transformation of compound **4**. Refluxing a solution of **4** with ceric ammonium nitrate (CAN) in acetonitrile for 6 hours afforded compound **2** in 70% yield. The oxidative transformation was clearly evident from the disappearance of aliphatic proton signals in the ¹H NMR spectrum of **4** at δ ∼2.4–2.6 ppm and the emergence of a downfield aromatic singlet at δ 6.50 ppm. Further The ¹³C NMR spectrum confirmed the conversion by the appearance of two carbonyl signals at δ 183.48 and 180.39 ppm, characteristic of the 1,4-dioxo-carbazole core.

Finally, the desired compound 6-methyl-9*H*-carbazole-3-carboxylic acid (**3**) was prepared through a two-step transformation of compound **4**. Initially, a Wolff–Kishner reduction was performed using hydrazine hydrate in ethylene glycol, which was followed by aromatization employing palladised charcoal to furnish carbazole **3** (Scheme 2). The transformation from **4** to **3** was supported by the upfield shift of the ¹H NMR spectrum, showing a methyl singlet at δ 2.46 ppm and aromatic signals in the δ 7.2–8.7 ppm range, with complete loss of aliphatic methylene and methine resonances which was also supported by ¹³C NMR via the disappearance of aliphatic CH□ and CH signals (δ ∼29–47 ppm).

### Biological evaluations

#### Antibacterial sensitivity profile of Synthetic compounds against selected Bacteria

Three target compounds **1**, **2** and **3** were synthesized, and their structures and antibacterial activities are summarized in Table 1. The agar well diffusion assay revealed that all compounds exhibited notable antibacterial effects. The inhibition zone diameters ranged from 8.13 to 17.13 mm against the Gram-positive strains *Bacillus cereus* and *Staphylococcus aureus*, while for the Gram-negative strains *Escherichia coli* and *Salmonella Typhimurium*, the zones measured 8.4–13.13 mm. These results indicate that the synthesized compounds are more effective against Gram-positive bacteria than Gram-negative strains.

**Table 1:** Zone of Inhibition (mm) by compounds 1, 2, 3 and Tetracycline Against Selected Bacteria.

| Compounds | *S. aureus*<br>(MTCC87) | *B. cereus*<br>(MTCC430) | *E. coli*<br>(MTCC46) | *S. Typhimurium*<br>(MTCC733) |
| --- | --- | --- | --- | --- |
| 3-methyl-1-tosylhydrazono-2,3,4,9-tetrahydrocarbazole-6-carboxylic acid (**1**) | 11.20 ± 0.20 | 17.13 ± 0.15 | 13.13 ± 0.012 | 15 ± 0.20 |
| 3-methyl-1,4-carbazoloquinone-6-carboxylic acid (**2**) | 8.13 ± 0.15 | 14.97 ± 0.25 | 8.03 ± 0.15 | 12.03 ± 0.15 |
| 6-methyl-carbazole-3-carboxylic acid (**3**) | 13.83 ± 0.15 | 13.13 ± 0.15 | 10.03 ± 0.15 | 11.03 ± 0.15 |
| Tetracycline | 16.13 ± 0.15 | 9.17 ± 0.15 | 14.97 ± 0.25 | 14.77 ± 0.25 |

### Strains in Well Diffusion Assay

The minimum inhibitory concentration (MIC) analysis of compounds **1**, **2** and **3** against four bacterial strains revealed distinct differences in antibacterial potency displayed in Figure 4 and Table 2. Against Gram-positive bacteria *S. aureus* (MTCC87), **2** exhibited the lowest MIC value (48.42 µg/ml), followed by **1** (53.19 µg/ml) and **3** (61.89 µg/ml), indicating slightly higher efficacy of **2** in inhibiting this Gram-positive pathogen. In the case of *B. cereus* (MTCC430), **3** demonstrated exceptional potency with an MIC of 12.73 µg/ml, significantly outperforming **1** (133.89 µg/ml) and **2** (271.21 µg/ml). For the Gram-negative strain *E. coli* (MTCC46), **2** showed the greatest activity (168.01 µg/ml), followed by **1** (187.43 µg/ml), whereas **3** displayed markedly lower efficacy (293.33 µg/ml). Interestingly, against *S. Typhimurium* (MTCC733), **1** exhibited the strongest inhibitory effect (50.08 µg/ml), far surpassing **3** (459.67 µg/ml) and **2** (474.71 µg/ml). Overall, the data suggest that **3** is particularly effective against *B. cereus*, **2** shows broad but moderate activity with notable potency against *E. coli* and *S. aureus*, while **1** is the most potent against *S. Typhimurium*, highlighting the strain-specific antibacterial potential of each compound. Moreover, the observed antibacterial profiles are consistent with the results of molecular docking studies, supporting the experimental data obtained through agar well diffusion and MIC assays.

**Figure 4:**
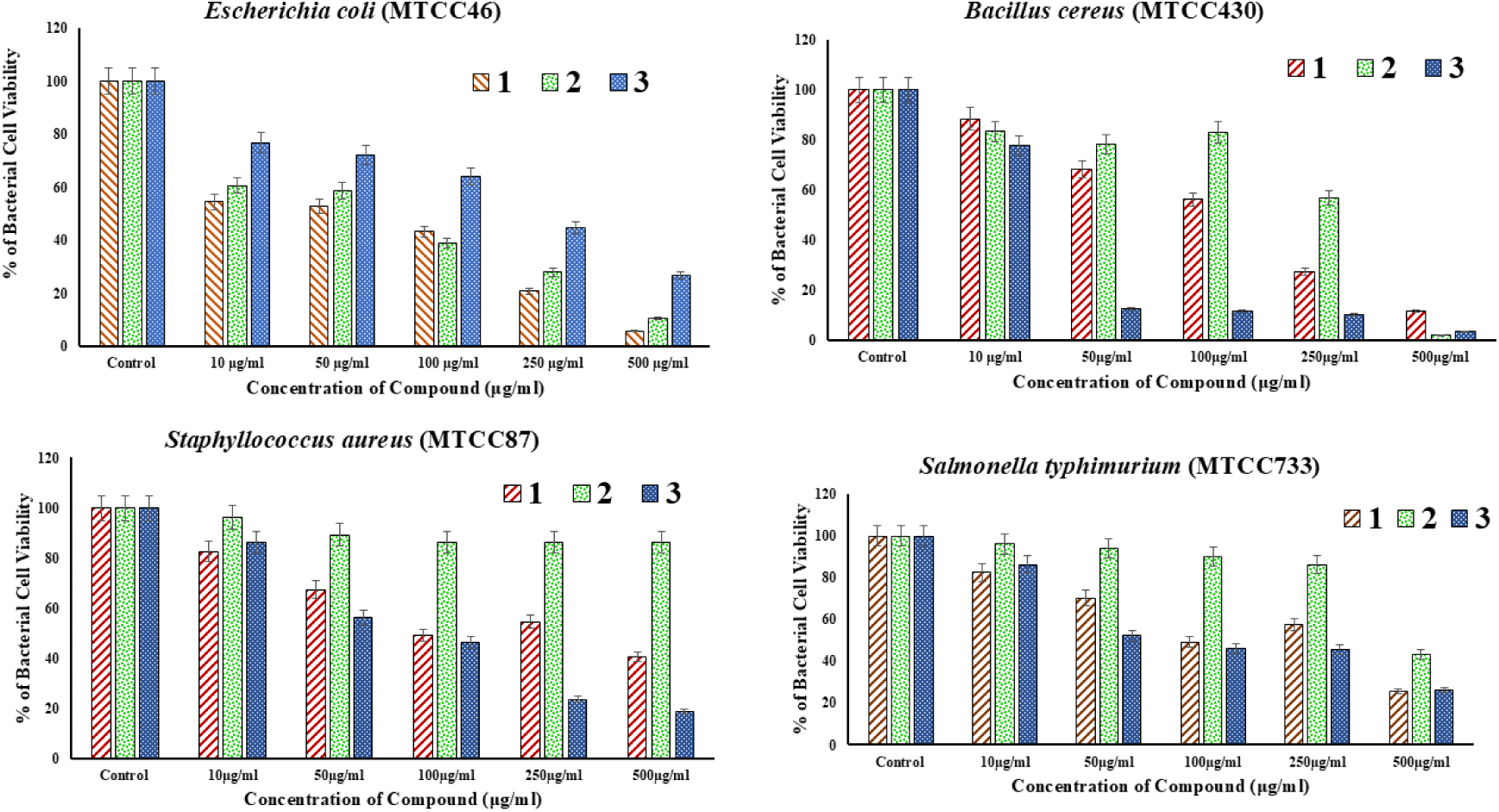
Dose response curves of *Escherichia coli* (A), *Salmonella Typhimurium* (B), *Bacillus cereus* (C) *and Staphylococcus aureus* (D) treated with hit compounds **1**, **2** and **3**. Bacterial survival was measured after overnight incubation with synthetic compounds to establish minimum inhibitory concentrations (MICs).

**Table 2:** Minimum Inhibitory Concentration (MIC) Values of compounds 1, 2 and 3 against selected Bacterial Strains.

| Compounds | *Staphylococcus aureus* (MTCC87) | *Bacillus cereus* (MTCC430) | *Escherichia coli* (MTCC46) | *Salmonella Typhimurium* (MTCC733) |
| --- | --- | --- | --- | --- |
| 3-methyl-1-tosylhydrazono-2,3,4,9-tetrahydrocarbazole-6-carboxylic acid (**1**) | 53.19 µg/ml | 133.89 µg/ml | 187.43 µg/ml | 50.08 µg/ml |
| 3-methyl-1,4-carbazoloquinone-6-carboxylic acid (**2**) | 48.42 µg/ml | 271.21 µg/ml | 168.01 µg/ml | 474.71 µg/ml |
| 6-methyl-carbazole-3-carboxylic acid (**3**) | 61.89 µg/ml | 12.73 µg/ml | 293.33 µg/ml | 459.67 µg/ml |

Among the three synthesized carbazole derivatives, 3-methyl-1,4-dioxo-4,9-dihydro-1*H*-carbazole-6-carboxylic acid (**2**) exhibited superior antibacterial activity compared to compounds **1** and **3**. Its more planar carbazole framework, combined with the presence of two keto groups at positions 1 and 4, likely contributes to enhanced interactions with bacterial targets. These electrophilic centers may facilitate covalent or strong non-covalent binding to nucleophilic residues in essential bacterial enzymes, thereby improving inhibitory potency. Additionally, the balance between hydrophobic and hydrophilic features in compound **2** may promote both adequate aqueous solubility and effective membrane penetration, further enhancing its antibacterial efficacy. This structural advantage underscores the potential of the 1,4-diketone carbazole motif as a promising lead for the development of potent antibacterial agents. These findings suggest that the 1,4-diketone-substituted carbazole framework represents a promising lead structure for the development of new antibacterial agents. In comparison with earlier studies in the field, the enhanced antibacterial activity of compound **2**, 3-methyl-1,4-dioxo-4,9-dihydro-1*H*-carbazole-6-carboxylic acid (**2**), stands out prominently. Prior investigations of carbazole-based antimicrobials, including derivatives with pyridinium moieties, carbazoloquinone alkaloids, and *N*-substituted analogs, have demonstrated only moderate to significant efficacy, with minimal inhibitory concentrations (MICs) often in the low microgram-per-milliliter range [40]. The structural novelty of compound **2** —the incorporation of dual keto functionalities into a planar carbazole core—introduces both electrophilic character and optimal physicochemical balance, achieving superior antibacterial potency compared to many of those previously reported scaffolds. This comparison underscores the critical role of the 1,4-diketone motif in potentiating carbazole derivatives as promising next-generation antibacterial agents.

### Molecular docking analysis

Computer-aided drug design (CADD) aims to support the discovery and development of drug candidates by reducing the overall cost and time associated with the drug design process. With advances in in silico methodologies and computational technologies, researchers can now perform more rapid and efficient optimization and identification of potential therapeutic agents. A key aspect of modern drug discovery involves structure-based computational modeling of ligand–receptor interactions. Among these approaches, molecular docking is widely utilized to predict the binding orientation and affinity of small molecules to their target macromolecules, such as proteins. Docking helps reveal conformational adaptations of both ligands and receptors in different environments and provides insights into molecular interactions at the atomic level. Although considered a relatively straightforward technique, molecular docking remains a powerful and effective tool in the early stages of drug design and was therefore chosen for the present study [41–43]. In the present study, molecular docking was employed to evaluate the binding affinity and interaction profile of the synthesized carbazole derivatives with biologically relevant protein targets.

### In vitro antibacterial activity Assessment

To complement the in vitro antibacterial studies, molecular docking was performed to investigate the binding interactions of the synthesized compounds **1, 2**, and **3** with selected bacterial gram +ve and gram -ve protein targets. This computational approach was employed to predict the binding affinity and possible binding modes of the compounds within the active sites of the target proteins. Docking simulations were conducted using an exhaustiveness level of 10 to ensure reliable sampling of conformational space. The key docking parameters, including binding energy scores and interaction profiles, are summarized in the Table 3.

**Table 3.**
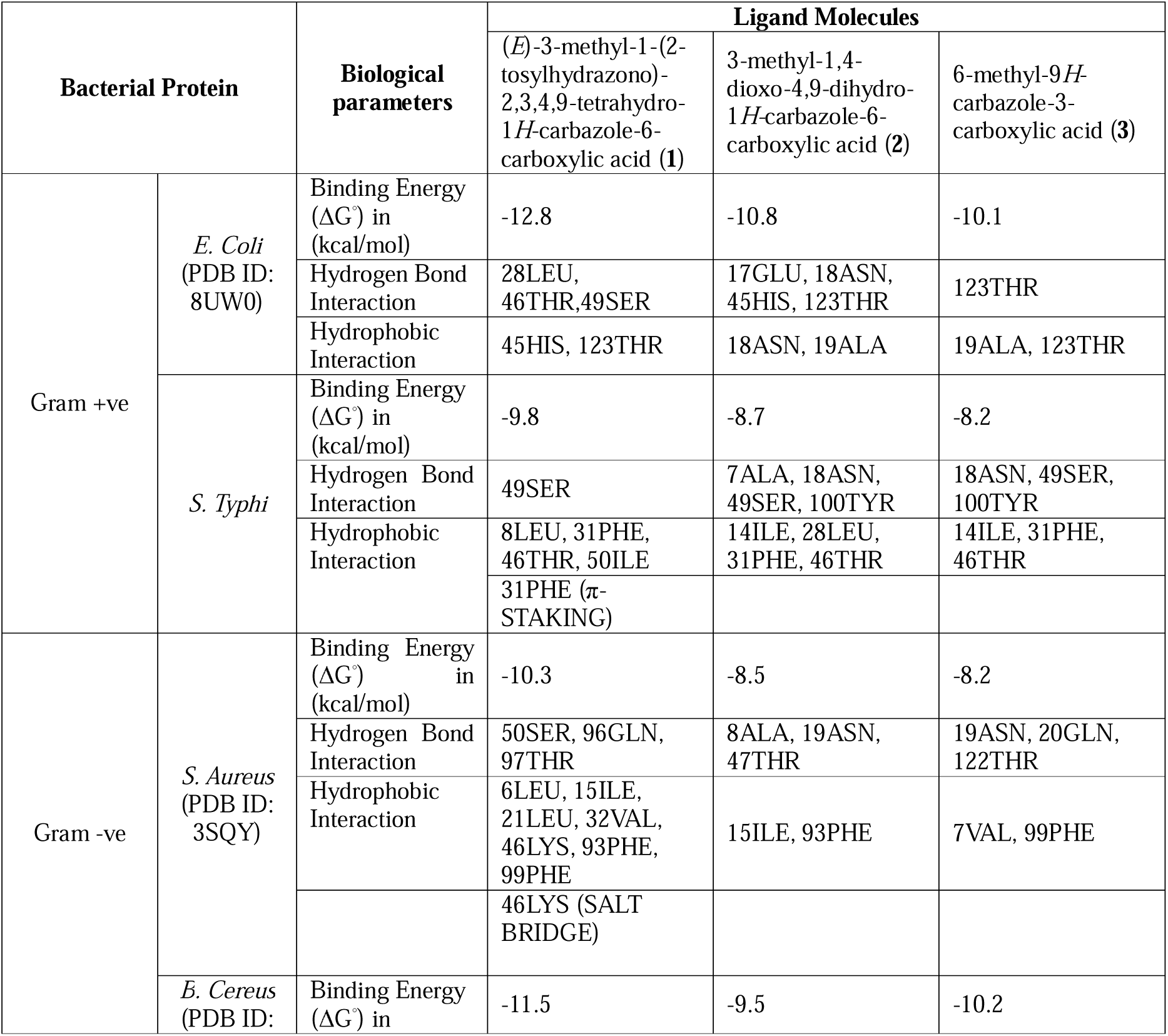
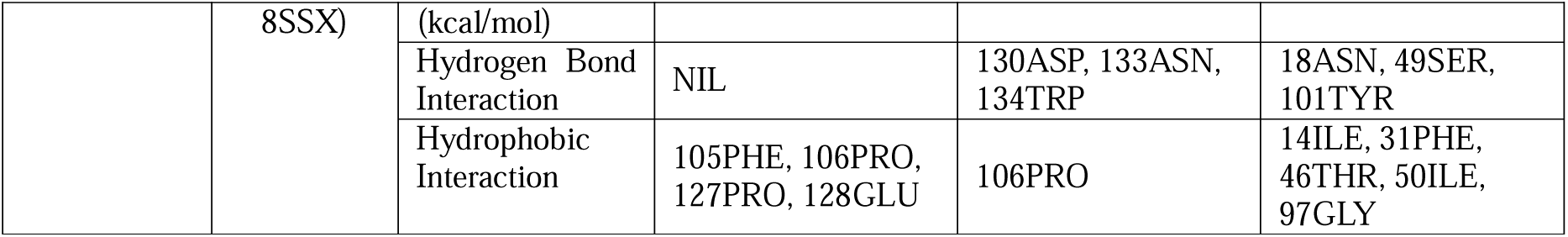
Molecular docking scores of compounds 1, 2 and 3.

Molecular docking analysis provided valuable insights into the role of specific atoms and functional groups within the synthesized ligands containing carbazole unit, that are critical for their interactions with microbial protein targets. By evaluating the binding affinities, the study highlighted the strength and specificity of these molecular interactions (Table 1) are the key factors in understanding the potential mechanisms of antibacterial action. The docking scores, expressed as free binding energies, reflect the stability and favourability of ligand–protein binding. Furthermore, the interaction profiles obtained were visualized using PyMOL, based on the “.pse” output files generated from PLIP analysis below.

Among the synthesized compounds, (*E*)-3-methyl-1-(2-tosylhydrazono)-2,3,4,9-tetrahydro-1*H*-carbazole-6-carboxylic acid (**1**) exhibited the highest binding affinities across all bacterial targets, with docking scores of –12.8 kcal/mol (*E. coli*), –9.8 kcal/mol (*S. typhi*), –10.3 kcal/mol (*S. aureus*), and –11.5 kcal/mol (*B. cereus*). In the E. coli DHFR active site, compound **1** showed a well-fitted binding pose, forming hydrogen bonds with residues LEU28, THR46, and SER49, along with hydrophobic interactions involving HIS45 and THR123. In S. typhi, key interactions included hydrogen bonding with SER49 and hydrophobic contacts with LEU8, PHE31, THR46, and ILE50, notably featuring π–π stacking with PHE31, which contributed significantly to binding stability. In the S. aureus DHFR pocket, hydrogen bonds were observed with SER50, GLN96, and THR97, while additional hydrophobic interactions were formed with LEU6, LEU21, LYS46, and PHE93. A noteworthy salt bridge interaction with LYS46 further enhanced the binding affinity. Although no hydrogen bonds were detected in the B. cereus complex (noted as "NIL"), the strong binding energy (–11.5 kcal/mol) was primarily driven by robust hydrophobic interactions with PHE105, PRO106, PRO127, and GLU128. These diverse interaction patterns—including hydrogen bonding, π-stacking, and salt bridge formation—highlight compound **1** as a promising broad-spectrum DHFR inhibitor with high binding stability across multiple bacterial targets.

**Figure 4.**
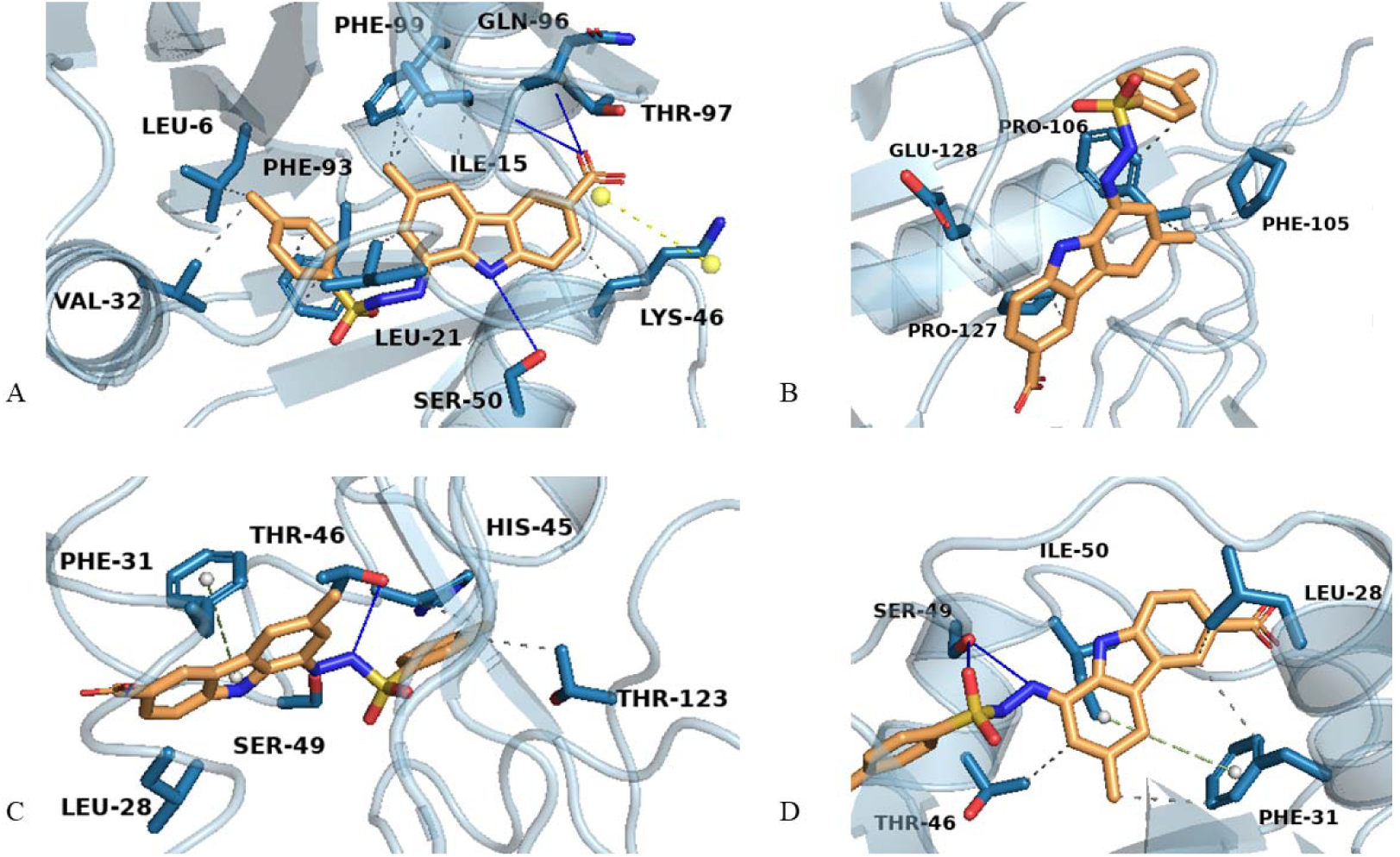
Ligand–protein interactions visualized through PyMOL, A: Compound **1** with 3SQY, B: Compound **1** with 8UVZ, C: Compound **1** with 8SSX, D: Compound **1** with S.Typhi

The 3-methyl-1,4-dioxo-4,9-dihydro-1*H*-carbazole-6-carboxylic acid (**2**) demonstrated consistent moderate-to-strong binding affinities across all bacterial DHFR targets, with docking scores of –10.8 kcal/mol (*E. coli*), –8.7 kcal/mol (*S. typhi*), –8.5 kcal/mol (*S. aureus*), and –9.5 kcal/mol (*B. cereus*). In *E. coli*, carbazoloquinone **2** formed multiple stabilizing hydrogen bonds with GLU17, ASN18, HIS45, and THR123, indicating a deeply anchored binding mode within the active pocket. Hydrophobic interactions with ASN18 and ALA19 further contributed to the stability of the complex. In *S. typhi*, a combination of hydrophobic contacts involving ILE14, LEU28, PHE31, and THR46, along with hydrogen bonds with ALA7, ASN18, SER49, and TYR100, supported a well-established binding orientation. For *S. aureus*, hydrophobic interactions with ILE15 and PHE93 complemented hydrogen bonds formed with ALA8, ASN19, and THR47, contributing to ligand stability. In the *B. cereus* DHFR complex, interactions were dominated by hydrophobic contacts with PRO106 and hydrogen bonds involving ASP130, ASN133, and TRP134. The widespread occurrence of hydrogen bonds across all complexes, along with adaptable hydrophobic interactions, underscores compound 2’s specificity and versatility as a DHFR-targeting agent. The key ligand–protein interactions of **2** are visualized through PyMOL using PLIP outputs are illustrated in Fig. 5.

**Figure 5.**
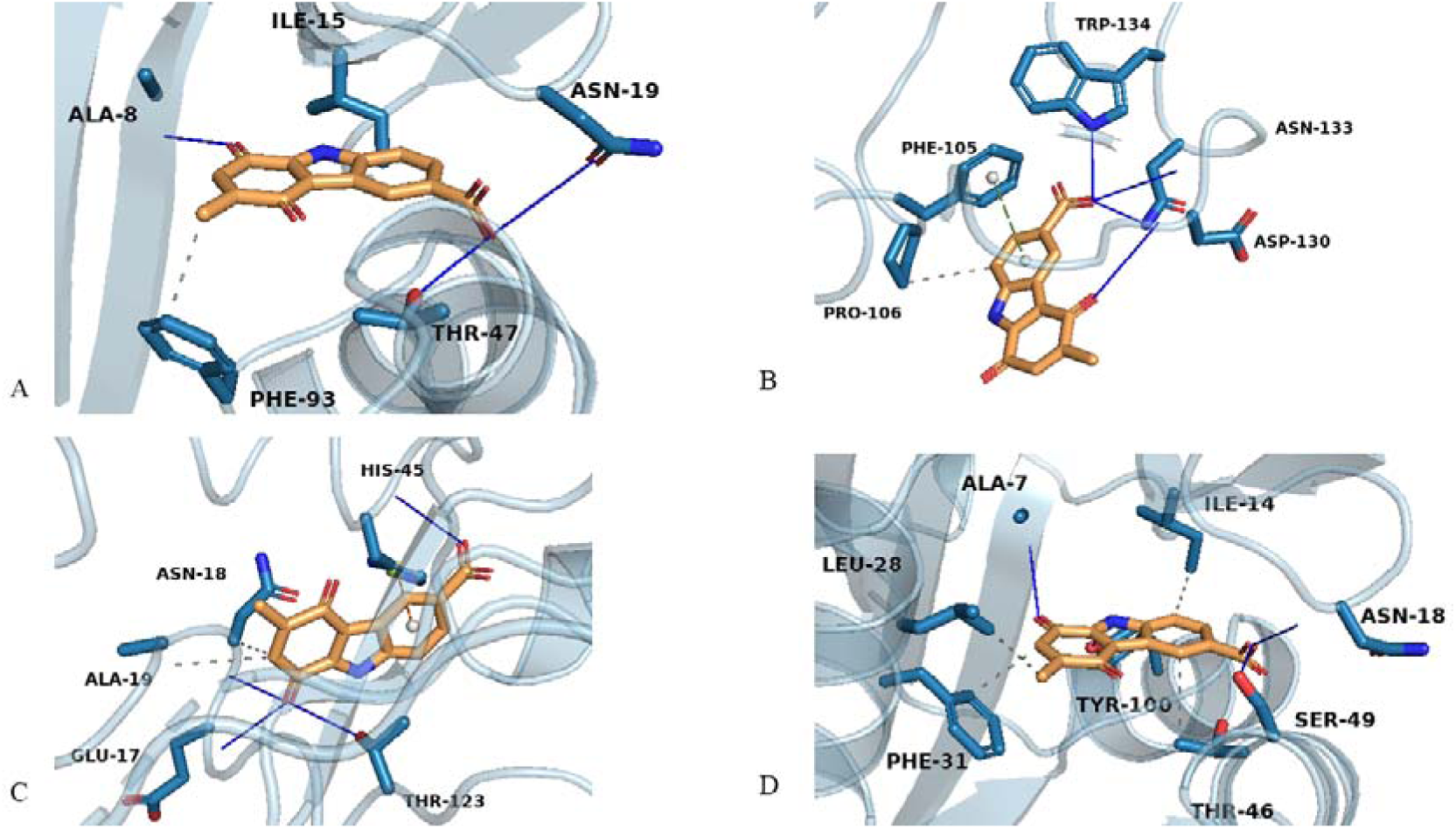
Ligand–protein interactions visualized through PyMOL. A: Compound **2** with 3SQY, B: Compound **2** with 8UVZ, C: Compound **2** with 8SSX, D: Compound **2** with S.Typhi

All four bacterial DHFR proteins exhibited moderate to strong binding affinities toward 6-methyl-9*H*-carbazole-3-carboxylic acid (**3**), with docking scores of –10.1 kcal/mol (*E. coli*), –8.2 kcal/mol *(S. typhi*), –8.2 kcal/mol (*S. aureus*), and –10.2 kcal/mol (*B. cereus*). Among them, the B. cereus DHFR complex demonstrated the most extensive ligand–protein contacts, suggesting species-specific binding optimization. In E. coli, the binding was characterized by hydrogen bonding with THR123 and hydrophobic interactions involving ALA19 and THR123, supporting a stable and localized binding mode. In S. typhi, a mix of polar and apolar interactions was observed, including hydrophobic contacts with ILE14, PHE31, and THR46, and hydrogen bonding with ASN18, SER49, and TYR100. The S. aureus complex featured hydrophobic interactions from VAL7 and PHE99, alongside hydrogen bonds with ASN19, GLN20, and THR122. A similar interaction profile was noted in B. cereus, where hydrophobic residues ILE14, PHE31, and THR46 anchored the ligand, while TYR101 and SER49 contributed to hydrogen bonding. These results highlight the carbazole scaffold as a versatile and reliable pharmacophore capable of maintaining binding fidelity across both Gram-positive and Gram-negative bacterial targets, with particularly strong affinity observed for B. cereus and E. coli. The ligand–protein interactions for compound **3** are recorded using PyMOL, as illustrated in Fig. 6.

**Figure 6.**
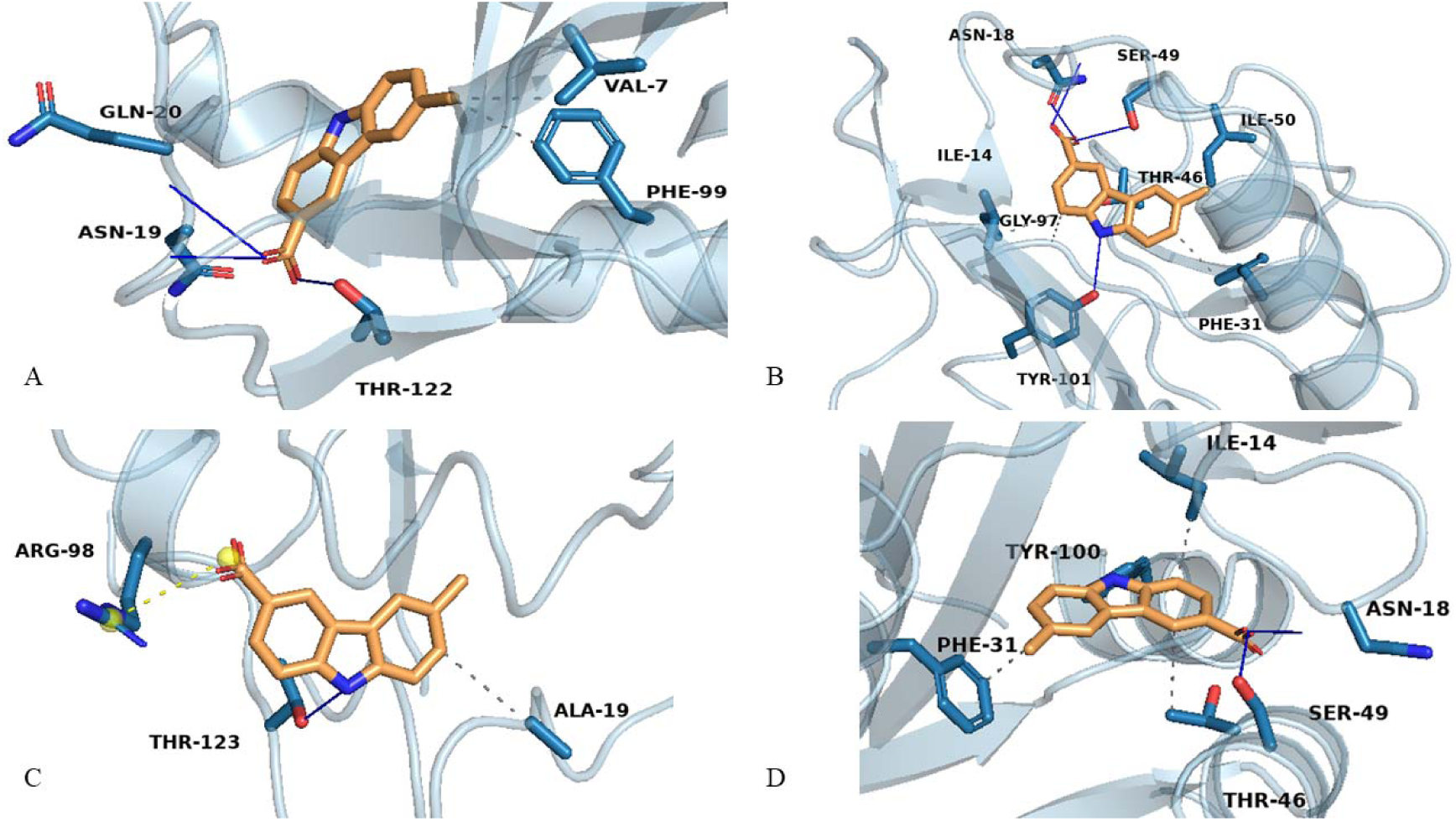
Ligand–protein interactions visualized through PyMOL. A: Compound **3** with 3SQY, B: Compound **3** with 8UVZ, C: Compound **3** with 8SSX, D: Compound **3** with S.Typhi

### Molecular Dynamic Simulations

The objective of the molecular dynamics (MD) simulation was to assess the structural stability and dynamic behaviour of the ligand–protein complexes over time. Upon binding, both the ligand and the protein often undergo conformational changes to achieve optimal fit within the active site. The resulting trajectories were analysed to extract key parameters, including the root mean square deviation (RMSD), which provides insights into the overall stability and compactness of the ligand– enzyme complexes throughout the simulation period. RMSD plots are particularly useful in monitoring structural fluctuations and confirming the robustness of the binding interactions over time [41].

In this case, to investigate the dynamic stability and structural adaptability of target proteins in response to ligand binding, Root Mean Square Deviation (RMSD) analysis was conducted for all systems over a 100 ns molecular dynamics simulation. RMSD provides insight into the overall deviation of protein backbone atoms from their initial conformation, reflecting conformational stability or flexibility induced by ligand interactions. Four bacterial proteins — *3SQY*, *8SSX*, *8UVZ*, and *Salmonella typhi DHFR (Typhi)* — were analyzed in both ligand-free (apoprotein) and ligand-bound forms with three candidate compounds **1**, **2** and **3**.

Among the three ligand–protein complexes analyzed, the 3SQY–Compound **1** system exhibited the highest RMSD deviation, reaching up to approximately 0.30 nm. (Fig. 7). This elevated value suggests that binding of the ligand **1** induces significant conformational changes and greater perturbations in the protein backbone, indicative of a more flexible or less stable interaction. The 3SQY–Compound **2** complex displayed moderately elevated RMSD values ranging between 0.18 and 0.25 nm. (Fig. 7), reflecting a moderate degree of conformational flexibility, yet remaining within a stable binding range. In contrast, the 3SQY–Compound **3** complex demonstrated the lowest fluctuations among the ligand-bound systems, with RMSD values ranging from 0.15 to 0.22 nm. (Fig. 7) and smooth trajectory patterns. This suggests that the ligand **3** fits well within the binding pocket, promoting stable accommodation with minimal disruption to the protein structure. The 3SQY apoprotein maintained a consistently low RMSD between 0.10 and 0.15 nm, confirming its high intrinsic structural stability in the absence of a ligand. Overall, these molecular dynamics results (Fig. 7) indicate that Carbazole **3** is the most compatible ligand for the 3SQY protein, supporting stable and tight binding with minimal conformational disturbance.

**Figure 7.**
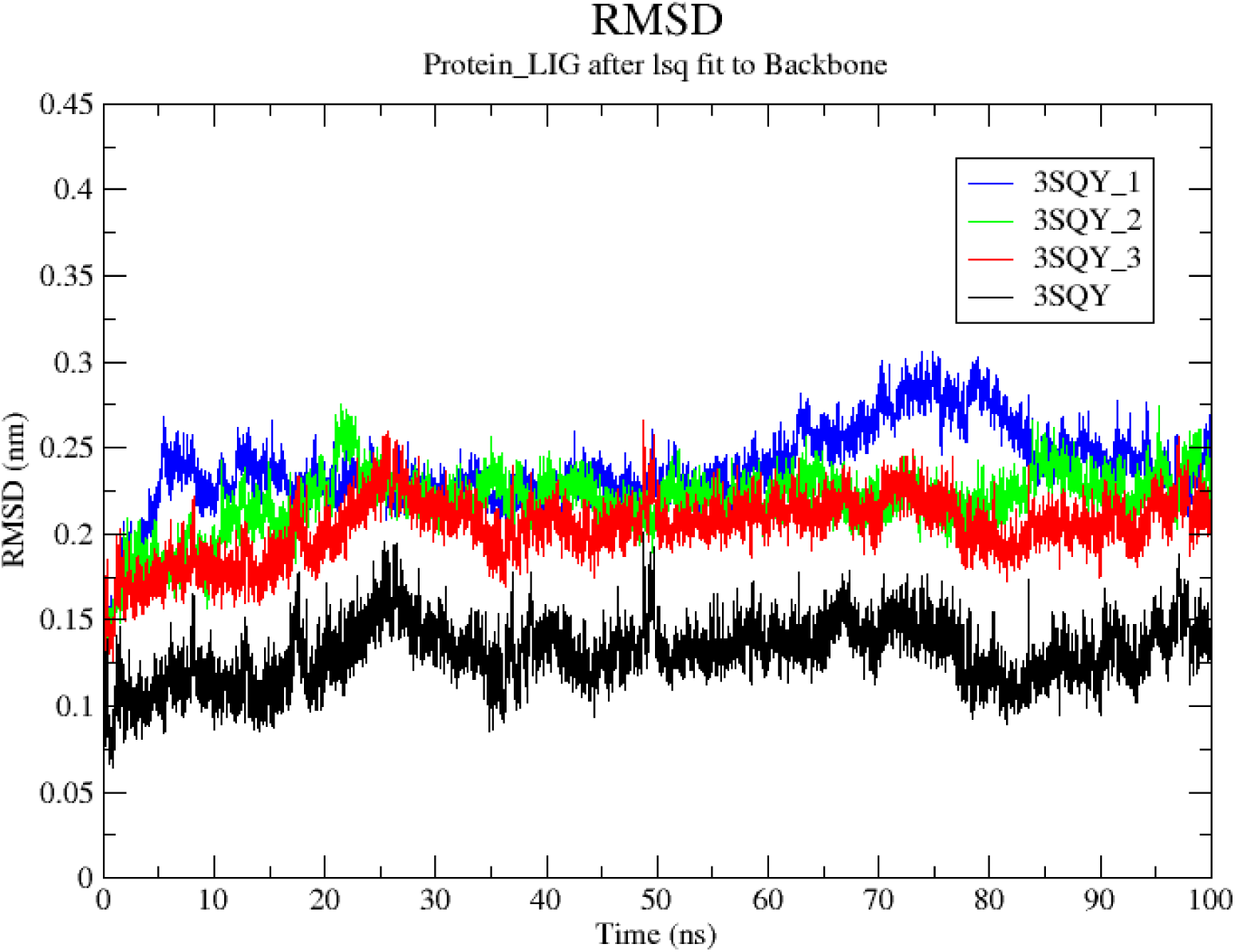
The RMSD/time plots presenting the molecular dynamics simulation trajectory of 3SQY with compound **1**, **2** and **3**: Black colour represents the 3SQY protein backbone, the blue colour represents compound **1**, the green colour represents compound **2** and the red colour represents compound **3**.

For the 8SSX apoprotein, RMSD values remained within the 0.10–0.20 nm range throughout the simulation, indicating a well-folded and intrinsically stable structure (Fig. 8). In contrast to the observations in the 3SQY system, Compound **1** binding to 8SSX resulted in the most stable complex, with RMSD values consistently ranging between 0.20 and 0.25 nm, suggesting minimal structural disruption and good compatibility. The 8SSX–Carbazoloquinone **2** complex demonstrated greater flexibility, with RMSD values rising to 0.35 nm. (Fig. 8) toward the later stages of the simulation, indicative of moderate conformational fluctuations. Notably, the 8SSX–6-methyl-carbazole-3-carboxylic acid **3** complex exhibited the highest RMSD, peaking around 0.40 nm, particularly after 60 ns of simulation time (Fig. 8). This suggests significant conformational changes and potential destabilization of the ligand–protein complex. This reversal in RMSD trends compared to the 3SQY system highlights the protein-specific nature of ligand compatibility, suggesting that **1** may be a more favorable and stabilizing ligand for 8SSX, whereas **3** demonstrates stronger compatibility with 3SQY.

**Figure 8.**
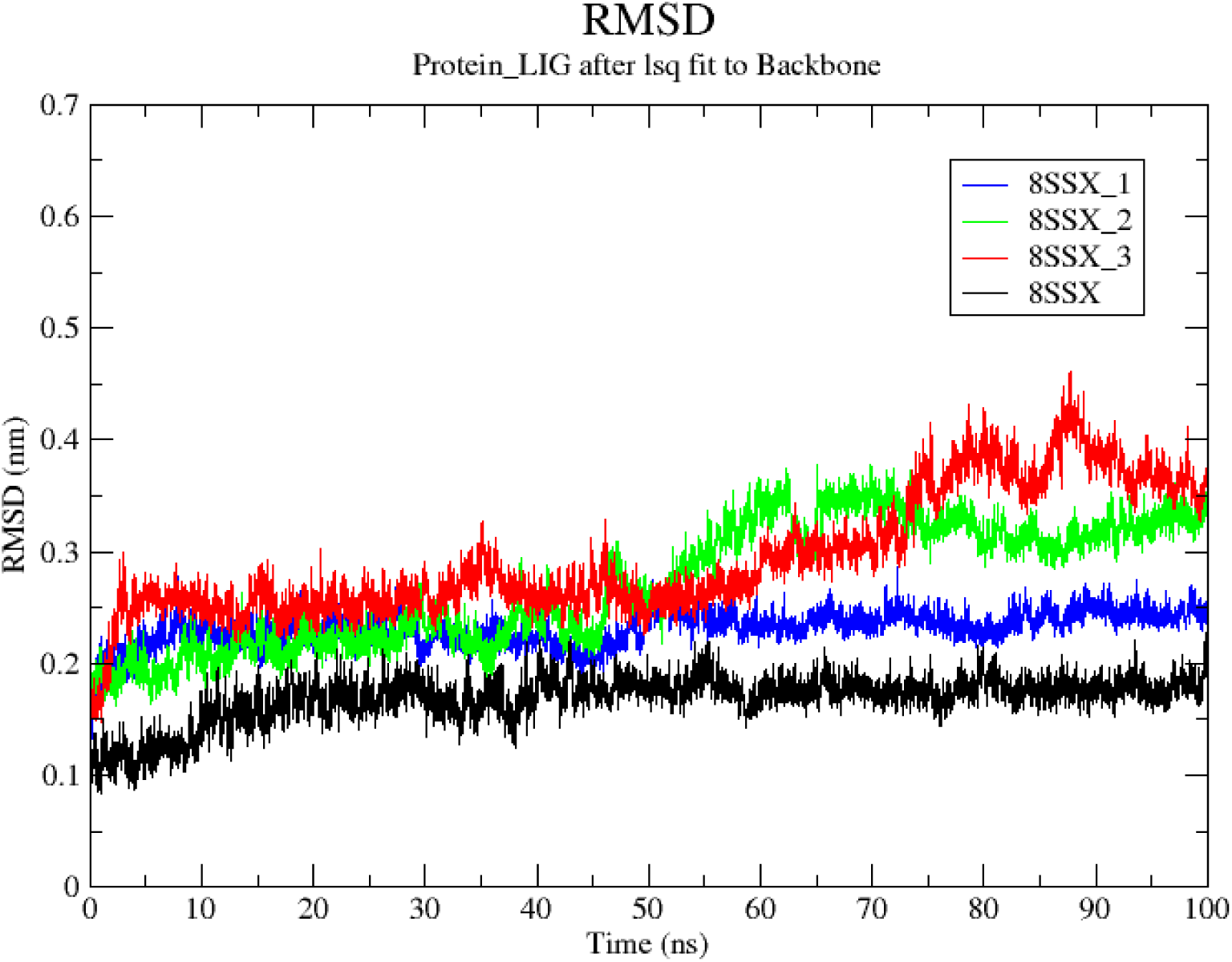
The RMSD/time plots presenting the molecular dynamics simulation trajectory of 8SSX with compound **1**, **2** and **3**: Black colour represents the 8SSX protein backbone, the blue colour represents compound **1**, the green colour represents compound **2** and the red colour represents compound **3**.

Again, the 8UVZ apoprotein maintained RMSD values between 0.15 and 0.20 nm, indicating a structurally consistent and stable conformation over the course of the simulation (Fig. 9). Both ligand-bound complexes, 8UVZ–Compound **1** and 8UVZ–Compound **3**, also exhibited relatively stable behavior, with RMSD values fluctuating modestly between 0.20 and 0.30 nm, suggesting good compatibility and minimal disturbance to the protein structure. In stark contrast, the 8UVZ– Compound **2** complex displayed pronounced structural instability, with multiple sharp RMSD spikes exceeding 0.60 nm (Fig. 9)—notably around 40 ns, 55 ns, and again toward the end of the simulation. These significant fluctuations imply that quinone binding induces major conformational changes or possible local unfolding events within the protein, undermining complex stability. Therefore, among the three ligands, Carbazoloquinone **2** appears to be the least compatible with 8UVZ, while 3-methyl-1-tosylhydrazonotetrahydrocarbazole-6-carboxylic acid (**1**) and 6-methyl-9*H*-carbazole-3-carboxylic acid (**3**) preserve structural integrity and demonstrate favourable dynamic behaviour.

**Figure 9.**
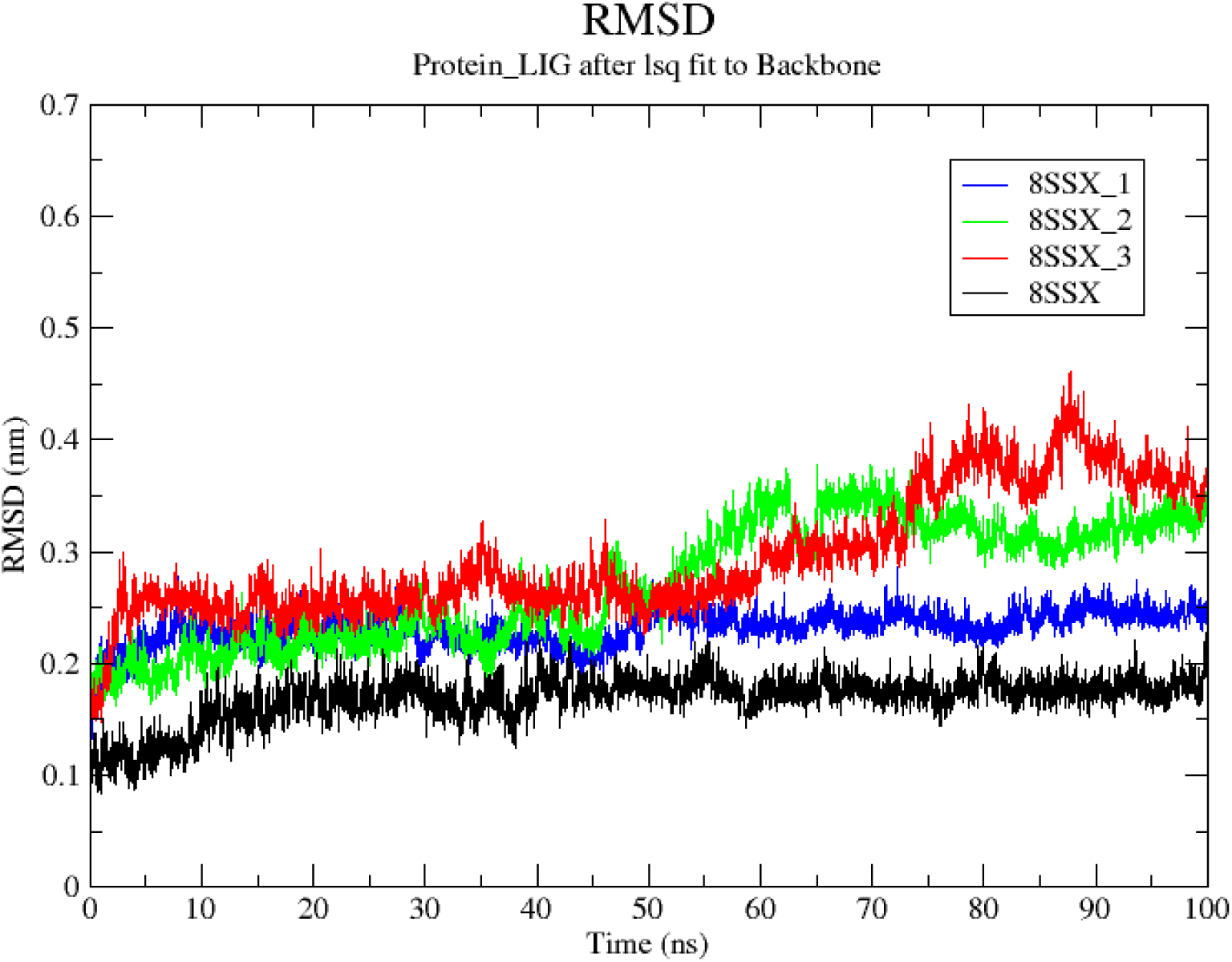
The RMSD/time plots presenting the molecular dynamics simulation trajectory of 8UVZ with compound **1**, **2** and **3**: Black colour represents the 8UVZ protein backbone, the blue colour represents compound **1**, the green colour represents compound **2** and the red colour represents compound **3**.

The RMSD profile of the *Salmonella typhi* DHFR apoprotein remained highly stable throughout the simulation, with deviations consistently within the 0.10–0.15 nm range, reflecting strong intrinsic structural integrity. Among the ligand-bound complexes, **1** again demonstrated the most stable behavior, maintaining RMSD values between 0.15 and 0.22 nm. (Fig. 10). The Compound **2**-bound complex exhibited slightly higher, yet acceptable RMSD values in the range of 0.20–0.32 nm, suggesting a degree of conformational flexibility while preserving overall complex stability. In contrast, the *S.Typhi*–Compound **3** complex showed notable instability, with RMSD values frequently spiking to approximately 0.50 nm between 30 and 60 ns before eventually stabilizing. This pattern suggests that **3** binding initially perturbs the protein’s structural integrity, compromising complex stability during the early simulation phase.

**Figure 10.**
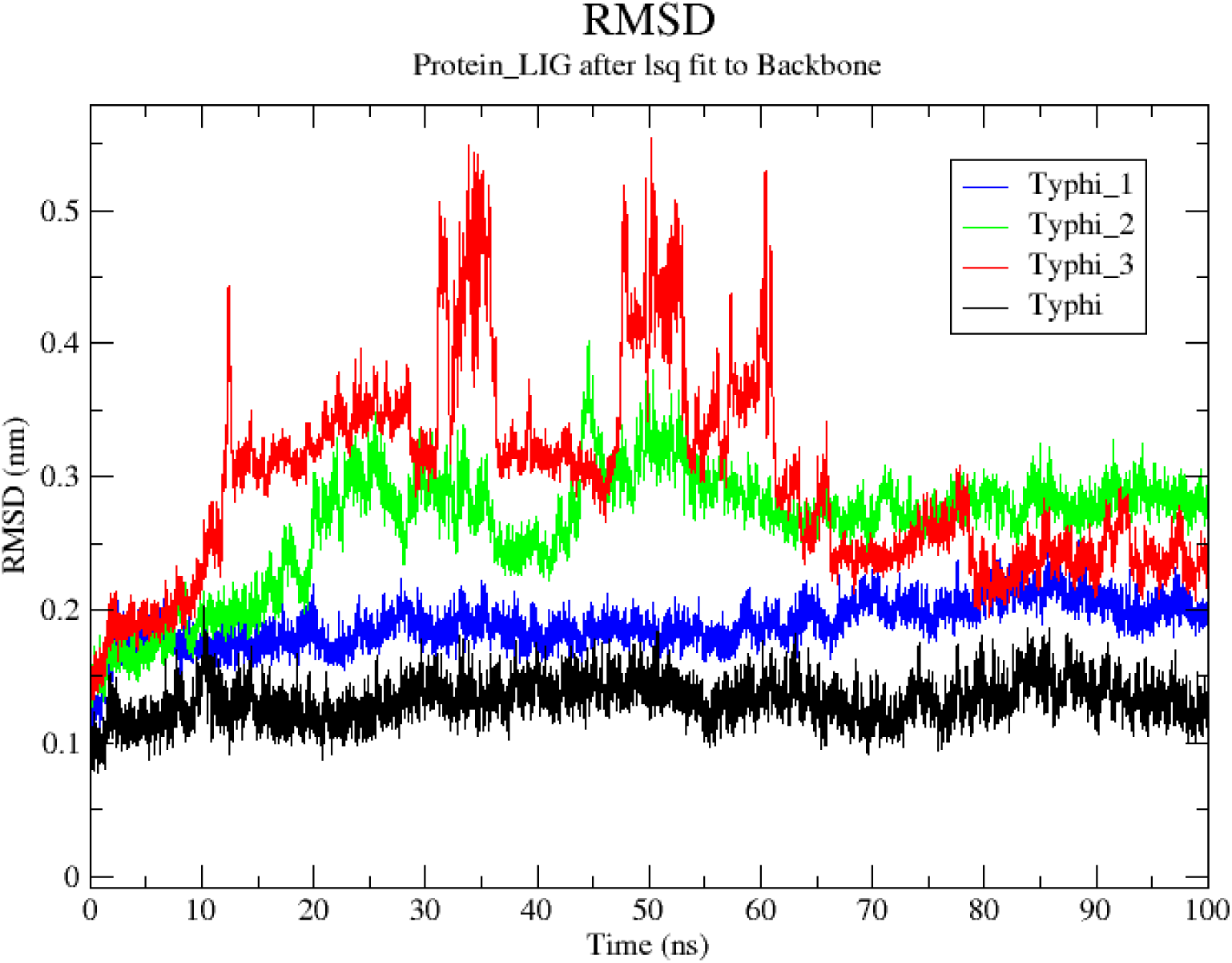
The RMSD/time plots presenting the molecular dynamics simulation trajectory of Salmonella typhi with compound **1**, **2** and **3**: Black colour represents the Salmonella typhi protein backbone, the blue colour represents compound **1**, the green colour represents compound **2** and the red colour represents compound **3**.

The molecular dynamics simulations of the ligand–protein complexes across four bacterial DHFR proteins—3SQY, 8SSX, 8UVZ, and *S. typhi* DHFR—revealed distinct patterns of structural stability and conformational response that are both ligand-dependent and protein-specific.

- 3-methyl-1-tosylhydrazonotetrahydrocarbazole-6-carboxylic acid (**1**) consistently exhibited low to moderate RMSD values across all protein systems, suggesting it is the most universally stable ligand. This uniform behavior points to a broad conformational tolerance by the DHFR active sites.
- 3-methyl-1,4-dioxo-4,9-dihydro-1*H*-carbazole-6-carboxylic acid (**2**) showed moderate dynamic stability with 3SQY, 8SSX, and *S. typhi*, indicating reasonably favourable interactions. However, it was markedly unstable with 8UVZ, exhibiting multiple RMSD spikes exceeding 0.6 nm. This suggests significant conformational strain and possible unfolding events.
- 6-methyl-9*H*-carbazole-3-carboxylic acid (**3**), while exhibiting excellent stability and tight binding with 3SQY, induced significant structural perturbations in both 8SSX and *S. typhi*. Its performance with 8UVZ was also suboptimal, with increased RMSD values indicating destabilization. These results emphasize the selective compatibility of compound **3**, underscoring the importance of target-specific ligand design for effective bacterial DHFR inhibition.

Overall, the RMSD analyses affirm the necessity of target-specific ligand optimization in rational drug design. Carbazole **3** may serve as a high-affinity scaffold for select targets like 3SQY, but its instability in other complexes limits its generalizability. In contrast, **1** has consistent stability across diverse DHFR variants, despite potentially lower binding specificity, suggests its suitability as a versatile core structure for developing broad-spectrum DHFR inhibitors. These structural insights will be critical for guiding docking validations, MM-PBSA free energy calculations, and the design of next-generation antibacterial agents against resistant strains.

### ADMET Analysis

To assess the drug-likeness and pharmacokinetic potential of the synthesized compounds, an in silico ADMET (Absorption, Distribution, Metabolism, Excretion, and Toxicity) analysis was carried out using the SwissADME web tool. The three acid-functionalized carbazole derivatives— compounds **1**, **2**, and **3**—were evaluated for key physicochemical and pharmacokinetic parameters, including molecular weight (MW), number of hydrogen bond donors (HBD) and acceptors (HBA), number of rotatable bonds, and fraction of sp³-hybridized carbons (Csp³). Additionally, each compound was subjected to multiple rule-based drug-likeness filters (Lipinski, Ghose, Veber, Egan, and Muegge) and alerts for potential substructures associated with non-specific binding or toxicity were assessed using PAINS and Brenk filters, while oral bioavailability scores and synthetic accessibility (SA) scores were also computed to estimate developability. A comparative overview of these parameters is presented through an ADMET radar plot (Fig. 11) and a heatmap visualization (Fig. 12), facilitating clear interpretation of the compounds’ drug-likeness profiles.

**Table 4.**
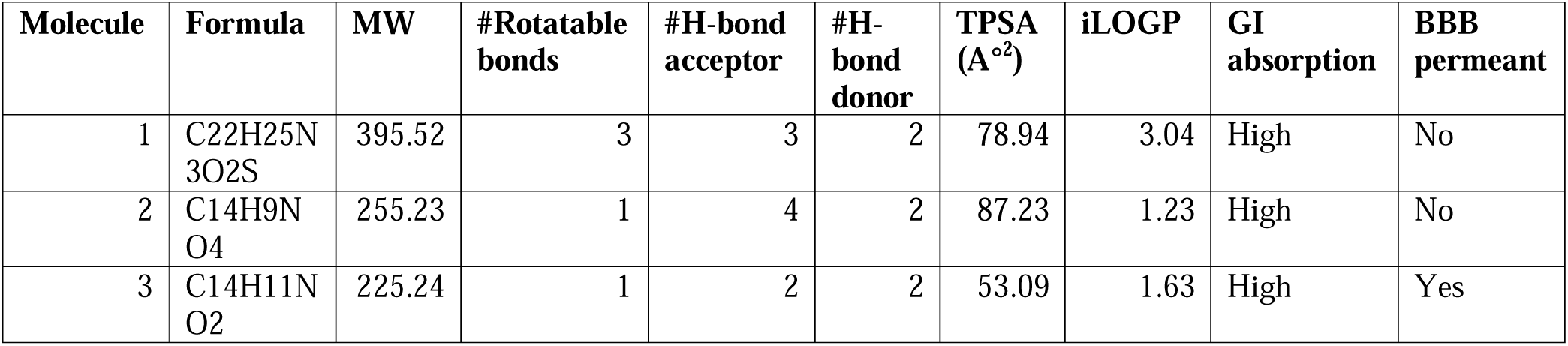
Druglikliness of compounds 1, 2 and 3 predicted by SWISSADME.

**Figure 11.**
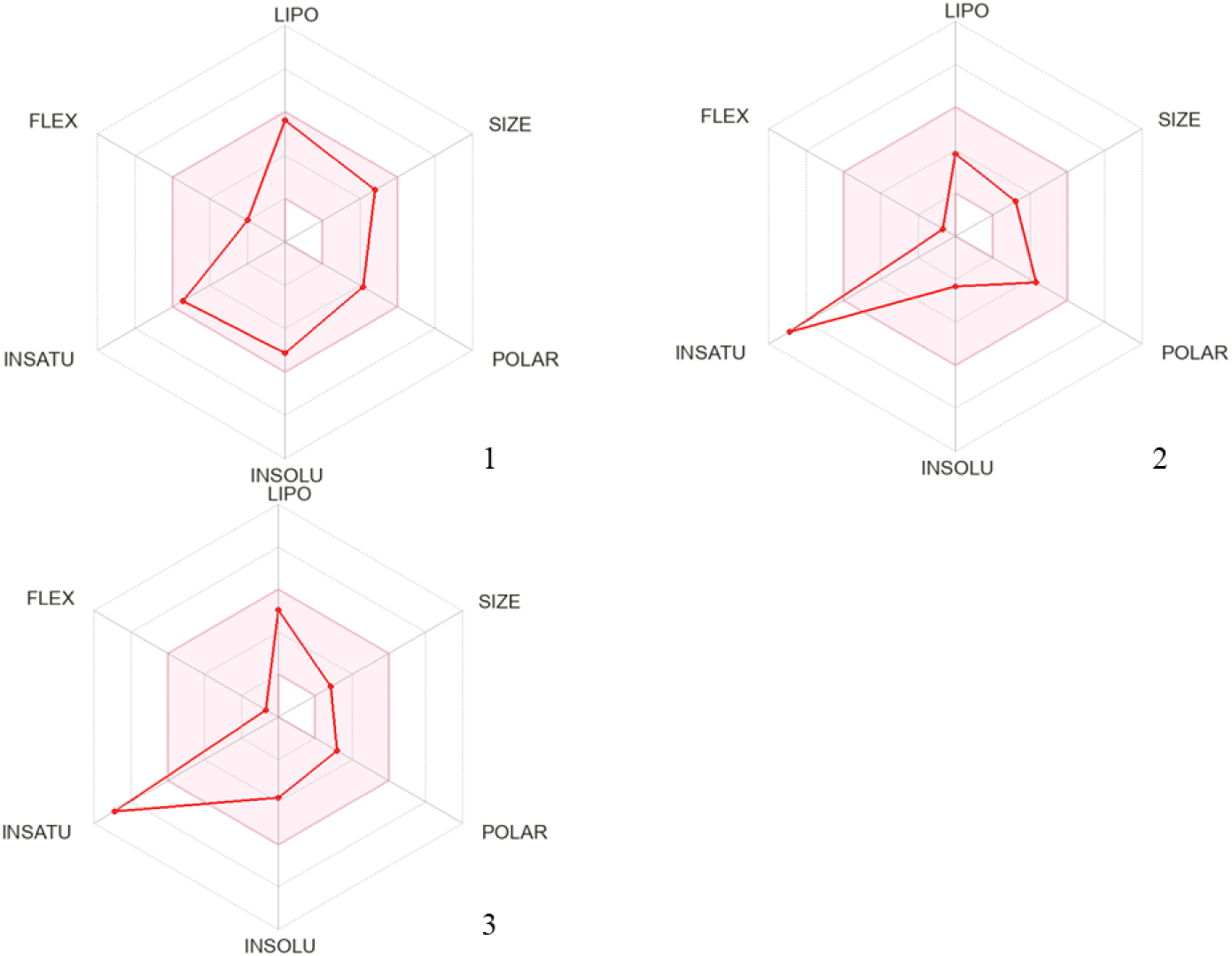
The bioavailability radars of **1**, **2** and **3** compounds

The compound **1** had the highest MW (395.52 g/mol) and synthetic complexity (score: 4.43), alongside two lead-likeness rule violations, one PAINS alert, and one Brenk alert. Although it had the highest Csp³ fraction (0.32), suggesting greater 3D molecular complexity, these advantages were offset by a relatively lower bioavailability score (0.55) and multiple alert flags. Its expanded radar profile (Fig. 11) and clustered heatmap values (Fig. 12) highlight these distinctions.

**Figure 12.**
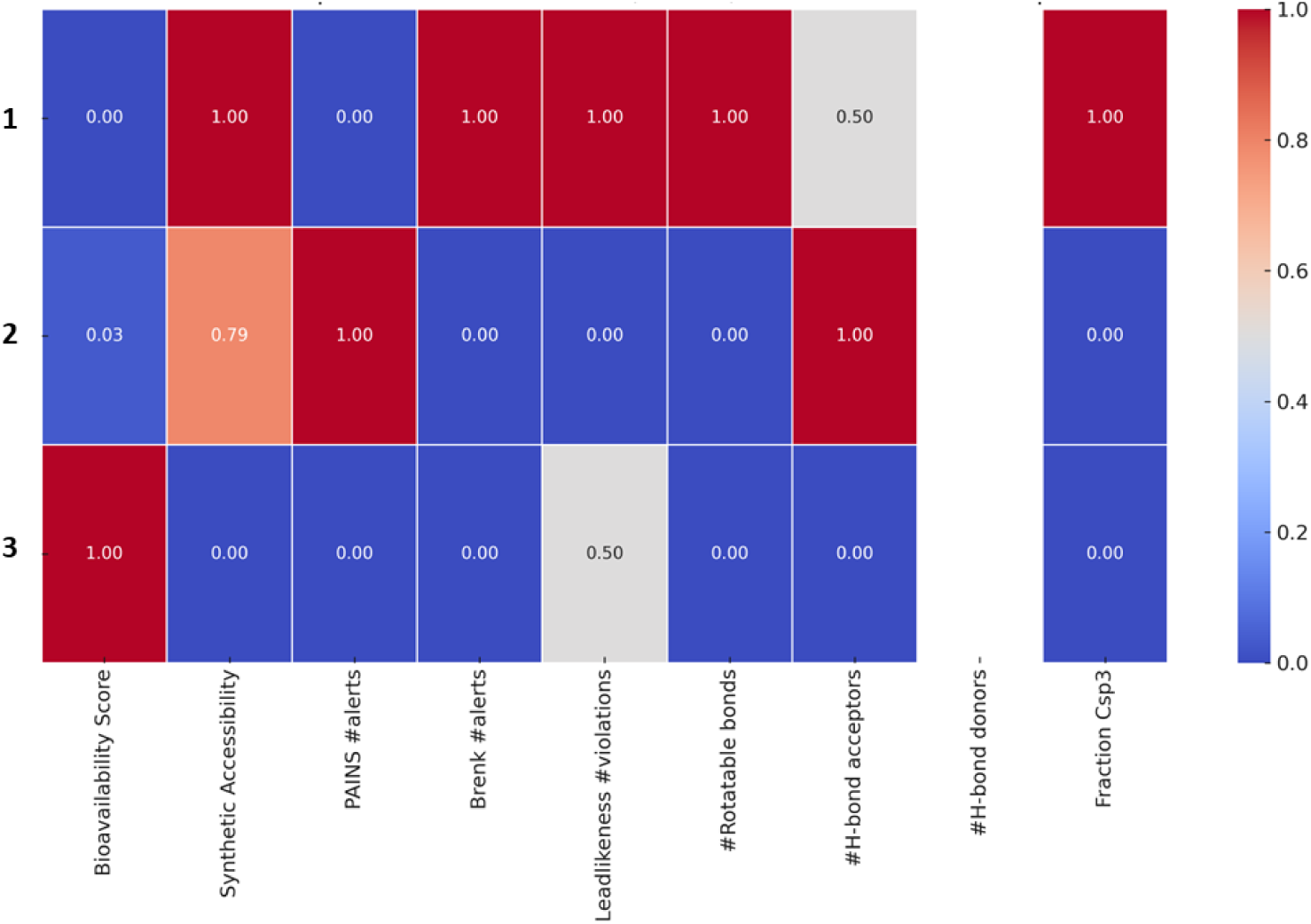
ADMET Heatmap representation of **1**, **2** and **3**

Carbazoloquinone acid derivative **2**, while slightly heavier (255.23 g/mol) and more synthetically complex (score: 3.78), also showed good compliance with drug-likeness criteria, with only a single PAINS alert noted. It had a higher number of hydrogen bond acceptors (4), suggesting better solubility characteristics, though its oral bioavailability score (0.56) was lower than that of Carbazole. In the radar plot (Fig. 11), Quinone’s profile shows slight deviations in certain parameters compared to Carbazole, consistent with the heatmap data (Fig. 12).

6- methyl-carbazole-3-carboxylic acid **3** exhibited the most favourable ADMET profile, with a low MW (225.24 g/mol), minimal synthetic complexity (synthetic accessibility score: 1.33), and the highest predicted oral bioavailability (0.85). It adhered strictly to all major drug-likeness rules and did not trigger any PAINS or Brenk structural alerts, indicating a chemically clean and pharmacologically viable scaffold. These favourable features are clearly represented in both the radar plot, where compound **3** occupies a balanced area across most axes (Fig. 11), and in the heatmap, which shows minimal alerts and violations (Fig. 12).

In summary, the ADMET data support 6-methyl-carbazole-3-carboxylic acid **3** as the most promising lead candidate, combining structural simplicity, high predicted oral bioavailability, and a clean alert profile.

## Conclusion

The present work successfully reports the design, synthesis, characterisation and comprehensive evaluation of a new class of acid-functionalized carbazole derivatives as potential antibacterial agents. The synthetic strategy successfully introduced acidic moieties into the carbazole scaffold, thereby enhancing physicochemical properties and enabling diverse biological interactions. Biological screening revealed that all three compounds exhibited measurable antibacterial activity, with distinct strain-specific potencies. Compound **2** emerged as the most promising derivative, displaying broad-spectrum efficacy with particularly strong activity against *S. aureus* and *E. coli*, supported by low MIC values. Compound **3** demonstrated remarkable selectivity against *B. cereus*, while compound **1** showed potent inhibition of *S. Typhimurium*, underscoring the role of structural modifications in modulating antibacterial profiles. Molecular docking studies complemented the biological results, revealing favourable binding affinities of all derivatives toward bacterial DHFR proteins, with compound **1** exhibiting the highest overall binding scores. Detailed interaction analyses highlighted the importance of hydrogen bonding, hydrophobic contacts, and π–π stacking in stabilizing ligand–protein complexes. Notably, the carbazoloquinone framework in compound **2** conferred superior planarity and electrophilic character, facilitating enhanced target engagement and suggesting its potential as a new lead scaffold for antibacterial drug development. Molecular dynamics simulations provided deeper insight into the structural adaptability and stability of the ligand–protein complexes. Compound **1** consistently demonstrated broad conformational compatibility across diverse DHFR proteins, suggesting its potential as a universal scaffold for broad-spectrum inhibition. In contrast, 6-methyl-carbazole-3-carboxylic acid **3** exhibited highly stable binding with 3SQY but induced destabilizing perturbations in other DHFR variants, indicating target-specific selectivity. Compound **2** showed intermediate stability but displayed marked incompatibility with 8UVZ, underscoring the importance of protein-specific optimization. Complementary ADMET analysis supported the pharmacokinetic viability of the synthesized derivatives. All substances met the parameters of Lipinski’s drug-likeness recommendations, according to pharmacokinetic studies. Among them, compound **3** demonstrated the most favourable profile, with high predicted oral bioavailability, minimal synthetic complexity, and no toxicity-related alerts, reinforcing its potential as a safe and pharmacologically tractable lead. Overall, this integrated experimental and computational study establishes acid-functionalized carbazole scaffolds as promising antibacterial candidates. These findings not only expand the chemical space of carbazole-based antimicrobials but also underscore the value of acidic functionalities in improving antibacterial efficacy. Future efforts will focus on structural optimization, in vivo validation, and exploration of synergistic activity to advance these carbazole derivatives toward the development of next-generation antibacterial agents capable of combating resistant pathogens.

## Experimental

### General Information

All the chemicals of reagent grade were purchased from several commercial sources and used without further purification. All reactions were performed under air and thin layer chromatography (TLC) was performed on Merck precoated TLC (silica gel 60 F_254_) plates. Melting points (°C) of the synthesised compounds were recorded in in open capillary tubes on a Digital melting point apparatus (Labard Scientific) and are uncorrected. The IR spectra were recorded in KBr discs on Schimadzu FTIR-8500. The ^1^H NMR (300 MHz) and ^13^C NMR (75 MHz) spectra were recorded on a Bruker Advance 300 spectrometer using CDCl_3_ or DMSO-d_6_ as the solvent. High-resolution mass spectra (HRMS) data were recorded on Qtof Micro YA263.

### Synthesis of compound 8

Compound **6** was synthesized according to the literature [44]. To a solution of **6** (2.2 g, 15.0 mmol) in methanol (20 mL), an aqueous solution of sodium acetate trihydrate (3.5 g) in water (15 mL) was added and allowed to cool at 0 °C. Then a solution of 4-carboxybenzenediazonium chloride (prepared from 2.15 g, 15.0 mmol 4-aminobenzoic acid) was added dropwise with stirring at 0 °C. After complete addition of dazonium solution the stirring was continued for 6 hrs at room temp. The yellow solid compound **8**, thus obtained, was collected by filtration and washed thoroughly with water. Upon crystallization from aqueous methanol, **8** (3.3 g) was obtained as yellow crystals

#### (*E*)-4-(2-(4-methyl-2-oxocyclohexylidene)hydrazinyl)benzoic acid (8)

Yellow solid; isolated yield: 86%, R_f_ = 0.6 (hexane:ethylacetate, 8:2); mp 274–276 °C; ¹H NMR (300 MHz, DMSO-d□, δ ppm): 13.37 (s, 1H, COOH), 10.17 (s, 1H, NH), 7.84 (d, J = 8.7 Hz, 2H, Ar–H), 35 (d, J = 8.7 Hz, 2H, Ar–H), 2.83–2.73 (m, 1H, CH□), 2.52–2.43 (m, 5H, CH□), 1.54–1.40 (m, 1H, CH), 0.96 (d, J = 8.7 Hz, 3H, CH□). ¹³C NMR (75 MHz, DMSO-d□, δ ppm): 195.05 (C=O), 167.64 (COOH), 148.71 (C), 140.85 (C), 131.32 (×2, CH), 123.31 (C), 114.00 (×2, CH), 48.73 (CH), 29.57 (CH□), 28.88 (CH□), 25.95 (CH□), 21.52 (CH□). HRMS (m/z) [M+H]^+^ calcd for C_14_H_17_N_2_O□: 261.1239, found: 261.1235.

### Synthesis of compound 4

A solution of **8** (4.1 g, 0.01 mol) in glacial acetic acid (16.0 mL) containing conc. hydrochloric acid (4 mL) was heated to reflux for 4 min. The reaction mixture was poured into ice-water (100 mL) and the solid thus obtained was collected by filtration, washed with water thoroughly and dried. The residue was subjected to flash chromatography (hexane/ethylacetate, 7:3) on silica gel to give a white solid of **4** (1.8 g).

#### 3-methyl-1-oxo-2,3,4,9-tetrahydro-1*H*-carbazole-6-carboxylic acid (4)

White solid; isolated yield: 74%, R_f_ = 0.5 (hexane:ethylacetate, 7:3); mp 275-277 °C; ¹H NMR (300 MHz, DMSO-d□, δ ppm): 11.85 (s, 1H, COOH), 8.29 (s, 1H, Ar–H), 7.84 (d, *J* = 9.0 Hz, 1H, Ar–H), 7.43 (d, *J* = 9.0 Hz, 1H, Ar–H), 2.60–2.48 (m, 4H, CH□), 2.45–2.34 (m, 1H, CH), 1.10 (d, *J* = 8.0 Hz, 3H, CH□). ¹³C NMR (75 MHz, DMSO-d□, δ ppm): 191.16 (C=O), 168.54 (COOH), 140.88(C), 132.83(C), 129.55(C), 127.37(C), 125.78(Ar-CH), 124.59(C), 122.83(Ar-CH), 113.27 (Ar-CH), 46.63 (CH), 32.92(CH□), 29.27 (CH□), 21.51 (CH□). HRMS (m/z) [M+H]^+^ calcd for C□□H□□NO□: 244.0968, found: 244.0965.

### Synthesis of compound 1

A mixture of **4** (2.45g, 0.01 mol) and tosylhydrazide (1.86g, 0.01 mol) was taken in methanol (25.0 mL) and heated under reflux for 5 hrs. The resulting solution was then cooled to room temperature and kept overnight at cold condition to afford yellow solid of **1** (3.8 g). It was washed with cold methanol and dried. The compound **4** was further purified by crystallisation from methanol.

#### (*E*)-3-methyl-1-(2-tosylhydrazono)-2,3,4,9-tetrahydro-1*H*-carbazole-6-carboxylic acid **(1):**

Pale yellow crystals; isolated yield: 92%, R_f_ = 0.6 (hexane:ethylacetate, 7:3); mp 279–281 °C; ¹H NMR (300 MHz, DMSO-d□, δ ppm): 11.86 (s, 1H, COOH), 11.19 (s, 1H, NH), 10.44 (s, 1H, NH), 8.29 (s, 1H, Ar–H), 7.90 (d, *J* = 9.0 Hz, 2H, Ar–H), 7.84 (d, *J* = 9.0 Hz, 2H, Ar–H), 7.43 (d, *J* = 9.0 Hz, 2H, Ar–H), 3.09 (dd, 2H, CH□), 2.61–2.53 (m, 2H, CH□), 2.49 (s, 3H, CH□–Ar), 2.38–2.31 (m, 1H, CH), 1.10 (d, *J* = 8.0 Hz, 3H, –CH□). ¹³C NMR (75 MHz, DMSO-d□, δ ppm): 191.16 (C=N), 168.54 (COOH), 140.88(C), 132.83(C), 130.16(C), 129.56×2(Ar-CH), 128.36(C), 127.37×2(Ar-CH), 125.78(C), 125.38(Ar-CH), 124.60(Ar-CH), 122.83×2(C), 113.27 (Ar-CH), 46.63 (CH), 32.92(CH□), 29.27 (CH□), 21.51(CH□), 18.95 (CH□). HRMS (m/z) [M+H]^+^ calcd for C_21_H_22_N_3_O_4_S: 412.1326, found: 412.1323.

### Synthesis of compound 2

A solution of ceric ammonium nitrate (13.2 g, 24.0 mmol) in anhydrous acetonitrile (25-30 mL) and compound **4** (1.0 g, 4.0 mmol) was refluxed for 4 hrs. The resulting solution was then cooled to room temperature and kept overnight at cold condition to obtain bright red crystals of **2** (870.0 mg). The cyrstals was then filtered and washed with cold MeOH.

#### 3-methyl-1,4-dioxo-4,9-dihydro-1*H*-carbazole-6-carboxylic acid (2)

Red solid; isolated yield: 86%, R_f_ = 0.4 (hexane:ethylacetate, 7:3); mp 260-262 °C; ¹H NMR (300 MHz, DMSO-d□, δ ppm): 12.93 (s, 1H, COOH), 12.86 (s, 1H, NH), 8.54 (s, 1H, Ar–H), 7.86 (d, J = 8.4 Hz, 1H, Ar–H), 7.52 (d, J = 8.4 Hz, 1H, Ar–H), 6.50 (s, 1H, Ar–H), 1.98 (s, 3H, CH□). ¹³C NMR (75 MHz, DMSO-d□, δ ppm): 183.48 (C=O), 180.39 (C=O), 168.11 (COOH), 148.54 (C), 140.12 (C), 137.75 (CH), 132.15 (C), 127.36 (C), 126.50 (C), 124.52 (C), 123.48 (CH), 116.55 (CH), 114.27 (CH), 16.09 (CH□). HRMS (m/z) [M+H]^+^ calcd for C_14_H_10_NO_4_: 256.0610, found: 256.0608.

### Synthesis of compound 3

Compound **4** (1.0 g, 4.0 mmol) in ethylene glycol (10 ml) was heated with hydrazine hydrate (80%, 3.6 ml) and KOH (1.6 g) at 190 °C for 1 h and then up to 210 °C under reflux for 3 h. The reaction mixture was poured into ice-water and acidified with conc. HCl under cold-condition to obtain a solid a compound. Then the aqueous solution was extracted with dichloromethane (3 × 15 ml). The combined organic layer was washed with brine solution and dried over solid Na_2_SO_4_. Then the solvent was distilled off when a residue of the corresponding tetrahydrocarbazole derivative was obtained. The residue was utilized for the next step without further purification. Crude tetrahydrocarbazole was taken in decalin (10 ml) and then Pd/C (l0%, 150 mg) was added. The mixture was then refluxed for 5 h. After reaction, Pd/C was removed by filtration. Then the filtered decalin solution was extracted with 5% NaOH (10.0 mL×3). The resulting alkaline solution was then acidified with conc. HCl to obtain the ppt. of desired product **3**. The residue was subjected to flash chromatography on silica gel G and eluted with hexane-ethylacetate (9:1) to afford required 6-carboxycarbazole **3** (270.0 mg).

#### 6-methyl-9*H*-carbazole-3-carboxylic acid (3)

White solid; isolated yield: 54%, R_f_ = 0.7 (hexane:ethylacetate, 7:3); mp 254–255 °C; ¹H NMR (300 MHz, DMSO-d□, δ ppm): 11.51 (s, 1H, COOH), 11.02 (s, 1H, NH), 8.71 (s, 1H, Ar–H), 8.01 (d, *J* = 8.1 Hz, 1H, Ar–H), 7.64 (d, *J* = 8.1 Hz, 1H, Ar–H), 7.49 (d, *J* = 8.1 Hz, 1H, Ar–H), 7.42 (d, *J* = 8.1 Hz, 1H, Ar–H), 7.26 (t, 1H, Ar–H), 2.46 (s, 3H, CH□). ¹³C NMR (75 MHz, DMSO-d□, δ ppm): 169.68 (COOH), 143.75 (C), 139.65 (C), 137.04 (C), 129.29(C), 127.85(C), 123.44(CH), 122.70(CH), 121.59 (CH), 120.76(C), 112.14 (CH), 111.74 (CH), 110.48 (CH), 22.66 (CH□). HRMS (m/z) [M+H]^+^ calcd for C_14_H_12_NO_2_: 226.0868, found: 226.0865.

### Biological methods

#### Bacterial Strains and Culture Conditions

For the antibacterial activity assessment, four reference strains have been selected, such as Staphylococcus aureus (MTCC87), Bacillus cereus (MTCC430) (Gram-positive), and Escherichia coli (MTCC46), and Salmonella Typhimurium (MTCC733) (Gram-negative). These strains have been chosen due to their clinical relevance and routine use in antimicrobial testing protocols. The bacterial cultures have been preserved at −20□°C in 20% (v/v) glycerol stocks until required for experimentation. For reactivation, each strain will be inoculated into nutrient broth (NB), composed of beef extract (5 g/L), peptone (10 g/L), and sodium chloride (5 g/L), adjusted to pH 7.0–7.2. An inoculum of 1 mL of each bacterium has been transferred into 20 mL of NB and incubated at 37□°C under shaking at 100 rpm. The optical density (OD) of the cultures will be monitored at 600 nm using a Shimadzu UV-1800 spectrophotometer to ensure that bacterial cells remain in the exponential growth phase for subsequent testing.

### Antibacterial Activity Evaluation Agar Well Diffusion Assay

The antimicrobial activity was assessed using the agar well diffusion method. Nutrient agar was employed for evaluating antibacterial activity. Freshly cultured human, bacterial, fungal, and shrimp pathogenic strains, prepared in broth form, served as the test organisms. These microbial strains were uniformly spread onto the surface of the respective agar media using sterile cotton swabs. Wells measuring 5 mm in diameter were then created in the agar plates with the help of a sterile cork borer. Different bacterial strains were introduced into the wells using sterile dropping pipettes. The plates were allowed to stand for 1 hour at room temperature to enable pre-incubation, followed by incubation at 37°C for 24 hours. After incubation, the diameter of the inhibition zones was measured to determine antimicrobial efficacy.

### Determination of Minimum Inhibitory Concentration (MIC)

The broth dilution method was used to determine the MICs [45]. The minimum concentration of test compounds (**1**, **2**, and **3**) that inhibits the growth of the test bacteria was determined by broth and agar dilution methods, following CLSI guidelines (CLSI 2019) [46]. A standardized suspension of bacteria at 2×106 CFU/ml in MHB containing test agents (0-500 µg/ml) was incubated in a shaker incubator (200 rpm) at 37°C for 24h. The lowest concentration of the tube without noticeable growth was recorded as the MIC; the MBC was determined using a standardized suspension (1.0 ml) of the test bacteria in MHB containing test agents at 0- to 4-fold the MIC, incubated overnight at 37 °C. The aliquots (0.1 ml) at hourly intervals were collected to measure the OD at 600 nm and plated to count the viable bacterial colonies.

### Computational details

The selection of target proteins was based on their significance as therapeutic targets and their critical roles in microbial pathogenicity. Specially an important bacterial enzyme called dihydrofolate reductase (DHFR) is required for the manufacture of tetrahydrofolate (THF), a vital cofactor for the building blocks of nucleic acids and some amino acids. These processes can be interfered with by blocking DHFR, which kills bacteria and makes DHFR a viable target for novel antibacterial medications [47]. Following the reported protocol [48] molecular docking was performed. Crystal structures of bacterial protein (PDB ID: 8SSX, 8UW0 [49], 3SQY[50]) were retrieved from RCSB Protein Data Bank (PDB) (https://www.rcsb.org/). The crystal structure of DHFR protein of *S.Typhi* was not available, so the sequence was extracted from NCBI (Accession No. - AGK65722.1) and modelled using Swiss Model [51]. The protein structure was validated from SAVESv6.1 (with an ERRAT [52] score of 96.59% and PROCHECK [53] with 0 errors and a passed Ramachandran plot). Due to novelty of the ligands synthesized, the 2D structures had to be created using ChemDraw software in mol format. Ligand molecules were prepared (addition of polar H-Atoms, addition of Gasteiger charges, generation of 3D coordinates, conversion into pdbqt format) using OpenBabel software [54]. Protein structure preparation (addition of polar H-Atoms, addition of Kollman charges, grid preparation, conversion into pdbqt format) was carried out in AutoDock [55]. Finally, using AutoDock Vina [56,57] and a shell script the entire process of docking was automated. The protein-ligand interaction was studied by uploading the complexes to the protein-ligand interaction profiler (PLIP) [58], an online web site. PyMOL was used to visualise the output ". pse" file from PLIP.

### Molecular Dynamic Simulations

GROMACS [59] was used for molecular dynamics simulation investigations. To generate the ligand topology, the Swiss Param (https://www.swissparam.ch/) [60] server was utilised. After uploading the ligand molecule’s MOL2 coordinates, the server supplied the ligand topology zip file. The salt type was NaCl; the force field was set to CHARMM27 [61]; the water model was TIP3; the apo-protein and complexes were set to cubic and dodecahedron box types, respectively; the energy minimisation steps were set to 50,000; and the NPT and NVT equilibration was performed at 300 K with a simulation time of 100 ns.

### ADMET Analysis

Toxicological traits and pharmacokinetic investigations are crucial factors in choosing possible medication choices. In lieu of clinical trials, computer techniques were created to evaluate a possible drug candidate’s bioactivity [62]. SwissADME (http://www.swissadme.ch/citing.php) [63] using smiles notation as an input tabulates the physiological, pharmacokinetic and pharmacological properties of the possible drug candidates.

## Supplementary Information

The online version contains supplementary material

## Acknowledgements

The authors, SC and AM gratefully acknowledges the authorities and management of The Bhawanipur Education Society College for their invaluable support and for providing the necessary facilities to carry out the research work

## CONFLICT OF INTEREST

The authors declare no conflict of interest, financial or otherwise.

## Authors’ contributions

Data collection and output processing – Aniruddhya Mukherjee, Suchandra Chakraborty, Ananya Das Mahapatra, Khushhali Menaria Pandey; Analysis and Interpretation – Aniruddhya Mukherjee, Suchandra Chakraborty, Ananya Das Mahapatra; Writing – Aniruddhya Mukherjee, Suchandra Chakraborty, Ananya Das Mahapatra; Critical Review – Aniruddhya Mukherjee, Khushhali Menaria Pandey, Suchandra Chakraborty, Ananya Das Mahapatra.

## Authors details

^1^Aniruddhya Mukherjee, The Bhawanipur Education Society College, Dept. of Chemistry, Kolkata-700020, West Bengal, India. ^2^Ananya Das Mahapatra, Brainware University, Dept. of Chemistry, Barasat, Kolkata-700125, West Bengal, India. ^3^Khushhali Menaria Pandey, Maulana Azad National Institute of Technology, Link Road Number 3, Near Kali Mata Mandir, Bhopal-462003, Madhya Pradesh, India. Aditya Maity^b^ ^1^Suchandra Chakraborty, The Bhawanipur Education Society College, Dept. of Chemistry, Kolkata-700020, West Bengal, India.

## Notes

### Competing Interest Statement

The authors have declared no competing interest.

## References

1. Wang, R., Yin, X., Zhang, Y., Yan, W. (2018). Design, synthesis and antimicrobial evaluation of propylene-tethered ciprofloxacin-isatin hybrids. Eur. J. Med. Chem. 156, 580–586. 10.1016/j.ejmech.2018.07.025

2. Hu, Y. Q., Zhang, S., Xu, Z., Lv, Z. S., Liu, M. L., Feng, L. S. (2017). 4-Quinolone hybrids and their antibacterial activities. Eur. J. Med. Chem. 141, 335–345. 10.1016/j.ejmech.2017.09.050

3. Saadon, K. E., Taha, N. M., Mahmoud, N. A., Elhagali, G. A., & Ragab, A. (2022). Synthesis, characterization, and in vitro antibacterial activity of some new pyridinone and pyrazole derivatives with some in silico ADME and molecular modeling study. J. Iran. Chem. Soc. 19(9), 3899–3917. 10.1007/s13738-022-02575-y

4. Bi, Y., Liu, X.-X., Zhang, H.-Y., Yang, X., Liu, Z.-Y., Lu, J., Lewis, P. J., Wang, C.-Z., Xu, J.-Y., Meng, Q.-G., Ma, C., & Yuan, C.-S. (2017). Synthesis and Antibacterial Evaluation of Novel 3-Substituted Ocotillol-Type Derivatives as Leads. Molecules, 22(4), 590. 10.3390/molecules22040590

5. Carrel, M., Perencevich, E. N., & David, M. Z. (2015). USA300 methicillin-resistant Staphylococcus aureus, United States, 2000–2013. EmeR. Infe. Dis., 21(11), 1973. 10.3201/eid2111.150452

6. Hvistendahl, M. (2012). China Takes Aim at Rampant Antibiotic Resistance. Science, 336(6083), 795–795. 10.1126/science.336.6083.795

7. Yezli, S., & Li, H. (2012). Antibiotic resistance amongst healthcare-associated pathogens in China. Int. J. anti. age., 40(5), 389–397. 10.1016/j.ijantimicag.2012.07.009

8. Gao, F., Xiao, J., & Huang, G. (2019). Current scenario of tetrazole hybrids for antibacterial activity. Euro. J. med. Chem., 184, 111744. 10.1016/j.ejmech.2019.111744.

9. Fitzpatrick, M. C., Bauch, C. T., Townsend, J. P., & Galvani, A. P. (2019). Modelling microbial infection to address global health challenges. Nat. micro., 4(10), 1612–1619. 10.1038/s41564-019-0565-8.

10. Knölker, H. J., & Reddy, K. R. (2002). Isolation and synthesis of biologically active carbazole alkaloids. Chem. rev., 102(11), 4303–4428. 10.1021/cr020059j

11. Knolker, H. J. (2004). Transition metal complexes in organic synthesis, part 70#. Synthesis of biologically active carbazole alkaloids using organometallic chemistry. Curr. Org. Synt., 1(4), 309–331. 10.2174/1570179043366594

12. Bauer, I., & Knölker, H. J. (2011). Synthesis of pyrrole and carbazole alkaloids. Alka. synt., 203–253. 10.1007/128_2011_192

13. Schmidt, A. W., Reddy, K. R., & Kno□lker, H. J. (2012). Occurrence, biogenesis, and synthesis of biologically active carbazole alkaloids. Chem. rev., 112(6), 3193–3328. 10.1021/cr200447s

14. Chakraborty, D. P., Barman, B. K., & Bose, P. K. (1964). On the structure of girinimbine, a pyrano-carbazole derivative, isolated from Murraya koenigii Spreng.

15. Chakraborty, D. P., Barman, B. K., & Bose, P. K. (1965). On the constitution of murrayanine, a carbazole derivative isolated from Murraya koenigii Spreng. Tetrahedron, 21(2), 681–685. 10.1016/S0040-4020(01)82240-7

16. Das, K. C., Chakraborty, D. P., & Bose, P. K. (1965). Antifungal activity of some constituents of Murraya koenigii Spreng.

17. Zhang, F. F., Gan, L. L., & Zhou, C. H. (2010). Synthesis, antibacterial and antifungal activities of some carbazole derivatives. Bioo. Med. Chem. Let., 20(6), 1881–1884. 10.1016/j.bmcl.2010.01.159

18. Asche, C., Frank, W., Albert, A., & Kucklaender, U. (2005). Synthesis, antitumour activity and structure–activity relationships of 5H-benzo [b] carbazoles. Bioo. Med. Chem., 13(3), 819–837. 10.1016/j.bmc.2004.10.038

19. Bedford, R. B., & Betham, M. (2006). N-H Carbazole Synthesis from 2-Chloroanilines via Consecutive Amination and C− H Activation. J. Org. Chem., 71(25), 9403–9410. 10.1021/jo061749g

20. Bedford, R. B., Betham, M., Charmant, J. P., & Weeks, A. L. (2008). Intramolecular direct arylation in the synthesis of fluorinated carbazoles. Tetrahedron, 64(26), 6038–6050. 10.1016/j.tet.2008.01.143

21. Bombrun, A., & Casi, G. (2002). N-Alkylation of 1H-indoles and 9H-carbazoles with alcohols. Tet Let., 43(12), 2187–2190. 10.1016/S0040-4039(02)00217-4

22. Meragelman, K. M., McKee, T. C., & Boyd, M. R. (2000). Siamenol, a new carbazole alkaloid from Murraya siamensis. J. nat. pro., 63(3), 427–428. 10.1021/np990570g

23. Choi, T. A., Czerwonka, R., Fröhner, W., Krahl, M. P., Reddy, K. R., Franzblau, S. G., & Knölker, H. J. (2006). Synthesis and activity of carbazole derivatives against Mycobacterium tuberculosis. ChemMedChem: Chem. Ena. Dru. Disc., 1(8), 812–815. 10.1002/cmdc.200600002

24. Choi, T. A., Czerwonka, R., Forke, R., Jäger, A., Knöll, J., Krahl, M. P., Krause, T., Reddy, K. R., Franzblau, S. G., & Knölker, H.-J. (2008). Transition metals in organic synthesis-Part 83#: Synthesis and pharmacological potential of carbazoles. Medi. Chem. Res., 17(2), 374–385. 10.1007/s00044-007-9073-0

25. Sakano, K. (1980). New antibiotics, carbazomycins A and B. J. Anti., 33, 961–966. 10.7164/antibiotics.33.683

26. Hook, D. J., Yacobucci, J. J., O’Connor, S., Lee, M., Kerns, E., Krishnan, B., Matson, J., & Hesler, G. (1990). Identification of the inhibitory activity of carbazomycins B and C against 5-lipoxygenase, a new activity for these compounds. The J. Anti., 43(10), 1347–1348. 10.7164/antibiotics.43.1347

27. Kaneda, M., Naid, T., Kitahara, T., Nakamura, S., Hirata, T., & Suga, T. (1988). Carbazomycins G and H, novel carbazomycin-congeners containing a quinol moiety. J. Anti., 41(5), 602–608. 10.7164/antibiotics.41.602

28. Takeya, K., Itoigawa, M., & Furukawa, H. (1989). Triphasic inotropic response of guinea-pig papillary muscle to murrayaquinone-A isolated from Rutaceae. Euro. J. pharm., 169(1), 137–145. 10.1016/0014-2999(89)90825-X

29. Balamurali, R., & Prasad, K. R. (2001). Synthesis, characterization and pharmacological activities of 5, 6, 11, 12-tetrahydroindolo [2, 3-a] carbazole derivatives. Il Farmaco, 56(3), 229–232. 10.1016/S0014-827X(01)01035-7

30. Kahan, F. M., Kropp, H., Sundelof, J. G., & Birnbaum, J. (1983). Thienamycin: development of imipenem-cilastatin. J. Anti. Chemo., 12(suppl_D), 1-35. 10.1093/jac/12.suppl_D.1

31. Bush, K., & Bradford, P. A. (2016). β-Lactams and β-lactamase inhibitors: an overview. Col. Spr. Har. pers. med., 6(8), a025247. 10.1101/cshperspect.a025247

32. Drlica, K., & Zhao, X. (1997). DNA gyrase, topoisomerase IV, and the 4-quinolones. Micro. mole. bio. rev., 61(3), 377–392. 10.1128/mmbr.61.3.377-392.1997

33. Chakraborty, B., Chakraborty, S., & Saha, C. (2014). Antibacterial Activity of Murrayaquinone A and 6□Methoxy□3, 7□dimethyl□2, 3□dihydro□1H□carbazole□1, 4 (9H)□dione. Int. J. Micro., 2014(1), 540208. 10.1155/2014/540208

34. Saha, C., Chakraborty, A., & Chowdhury, B. K. (1996). A new synthesis of 4-deoxycarbazomycin B and its antimicrobial properties. Ind. J. chem. Sect. B: Organic chemistry, including medical chemistry, 35(7), 677–680.

35. Chakraborty, S., Chakraborty, B., Saha, A., Saha, C., Ghosh, T. K., & Bhattacharyya, I. (2017). Evaluation of antimicrobial activity of synthesized fluorocarbazole derivatives based on SAR. Ind. J. Chem., 56, 701–708. https://www.webofscience.com/wos/WOSCC/full-record/000432762200002

36. Chakraborty, B., Chakraborty, S., Bhattacharyya, I., & Saha, C. (2014). In Vitro Activity of Synthesized 6-Chlor-2-methyl-1H-carbazole-1, 4 (9H)-dione against Methicillin-Resistant Staphylococcus aureus. IOSR J. Appl. Chem, 7(11), 61–66.

37. C. Ainsworth, Org. Synth., Coll. Vol. IV 1963, 536–539.

38. C. Saha, A. Chakraborty, B. K. Chowdhury, Indian J. Chem., Sect. B: Org. Chem. Incl. Med. Chem. 1996, 35, 677–680.

39. Chakraborty, S., Chattopadhyay, G., & Saha, C. (2013). A tandem reduction–oxidation protocol for the conversion of 1□keto□1, 2, 3, 4□tetrahydrocarbazoles to carbazoles via tosylhydrazones through microwave assistance: efficient synthesis of glycozoline, clausenalene, glycozolicine, and deoxycarbazomycin B and the total synthesis of murrayafoline A. J. Het. Chem., 50(1), 91–98. 10.1002/jhet.999

40. Wang, P. Y., Fang, H. S., Shao, W. B., Zhou, J., Chen, Z., Song, B. A., & Yang, S. (2017). Synthesis and biological evaluation of pyridinium-functionalized carbazole derivatives as promising antibacterial agents. Bioo. medi. chem. let., 27(18), 4294–4297. 10.1016/j.bmcl.2017.08.040

41. Sousa, S. F., Fernandes, P. A., & Ramos, M. J. (2006). Protein–ligand docking: current status and future challenges. Pro.: struc., fun., bioinfo., 65(1), 15–26. 10.1002/prot.21082

42. Meng, X. Y., Zhang, H. X., Mezei, M., & Cui, M. (2011). Molecular docking: a powerful approach for structure-based drug discovery. Cur. Comp.-aid. Drug. Desi., 7(2), 146–157. 10.2174/157340911795677602

43. Chaudhari, R., Tan, Z., Huang, B., & Zhang, S. (2017). Computational polypharmacology: a new paradigm for drug discovery. Exp. opi. drug dis., 12(3), 279–291. 10.1080/17460441.2017.1280024

44. Chakraborty, S., & Saha, C. (2013). Total synthesis of carbazomycin G. Euro. J. Org. Chem., 2013(25), 5731–5736. 10.1002/ejoc.201300467

45. Barnes, L., Heithoff, D. M., Mahan, S. P., House, J. K., & Mahan, M. J. (2023). Antimicrobial susceptibility testing to evaluate minimum inhibitory concentration values of clinically relevant antibiotics. STAR prot., 4(3), 102512. 10.1016/j.xpro.2023.102512

46. Wayne, P. A. (2019). Clinical Laboratory Standards Institute. Performance Standards for Antimicrobial Susceptibility Testing 29^th^ Edition, No: 1, *CLSI Supplement* M 100, ISBN 978-1-68440-032-4. 19087, USA, 2019. 2019; 1–281.

47. He, J., Qiao, W., An, Q., Yang, T., & Luo, Y. (2020). Dihydrofolate reductase inhibitors for use as antimicrobial agents. Euro. J. Medi. Chem., 195, 112268. 10.1016/j.ejmech.2020.112268

48. Mukherjee, A., Pandey, K. M., Ojha, K. K., & Sahu, S. K. (2023). Identification of possible SARS-CoV-2 main protease inhibitors: in silico molecular docking and dynamic simulation studies. Beni-Suef Uni. J. Bas. App. Sci., 12(1), 69. 10.1186/s43088-023-00406-4

49. Hummel, J. R., Xiao, K.-J., Yang, J. C., Epling, L. B., Mukai, K., Ye, Q., Xu, M., Qian, D., Huo, L., Weber, M., Roman, V., Lo, Y., Drake, K., Stump, K., Covington, M., Kapilashrami, K., Zhang, G., Ye, M., Diamond, S., Yeleswaram, S., Macarron, R., Deller, M. C., Wee, S., Kim, S., Wang, X., Wu, L., & Yao, W. (2024). Discovery of (4-pyrazolyl)-2-aminopyrimidines as potent and selective Inhibitors of cyclin-dependent kinase 2. J. Medi. Chem., 67(4), 3112–3126. 10.1021/acs.jmedchem.3c02287

50. Li, X., Hilgers, M., Cunningham, M., Chen, Z., Trzoss, M., Zhang, J., Kohnen, L., Lam, T., Creighton, C., G. C., Nelson, K., Kwan, B., Stidham, M., Brown-Driver, V., Shaw, K. J., & Finn, J. (2011). Structure-based design of new DHFR-based antibacterial agents: 7-aryl-2, 4-diaminoquinazolines. Bioo. medi. chem. Let., 21(18), 5171–5176. 10.1016/j.bmcl.2011.07.059

51. Waterhouse, A., Bertoni, M., Bienert, S., Studer, G., Tauriello, G., Gumienny, R., Heer, F. T., de Beer, T. A. P., Rempfer, C., Bordoli, L., Lepore, R., & Schwede, T. (2018). SWISS-MODEL: homology modelling of protein structures and complexes. Nucl. aci. res., 46(W1), W296–W303. 10.1093/nar/gky427

52. Colovos, C., & Yeates, T. O. (1993). Verification of protein structures: patterns of nonbonded atomic interactions. Pro. sci., 2(9), 1511–1519. 10.1002/pro.5560020916

53. Laskowski, R. A., MacArthur, M. W., Moss, D. S., & Thornton, J. M. (1993). PROCHECK: a program to check the stereochemical quality of protein structures. App. Crys., 26(2), 283–291. 10.1107/s0021889892009944

54. O’Boyle, N. M., Banck, M., James, C. A., Morley, C., Vandermeersch, T., & Hutchison, G. R. (2011). Open Babel: An open chemical toolbox. J. chemin., 3(1), 33. 10.1186/1758-2946-3-33

55. Morris, G. M., Huey, R., Lindstrom, W., Sanner, M. F., Belew, R. K., Goodsell, D. S., & Olson, A. J. (2009). AutoDock4 and AutoDockTools4: Automated docking with selective receptor flexibility. J. comp. chem., 30(16), 2785–2791. 10.1002/jcc.21256

56. Eberhardt, J., Santos-Martins, D., Tillack, A. F., & Forli, S. (2021). AutoDock Vina 1.2. 0: new docking methods, expanded force field, and python bindings. J. chem. info. mod., 61(8), 3891–3898. 10.1021/acs.jcim.1c00203

57. Trott, O., & Olson, A. J. (2010). AutoDock Vina: improving the speed and accuracy of docking with a new scoring function, efficient optimization, and multithreading. J. comp. chem., 31(2), 455–461. 10.1002/jcc.21334

58. Adasme, M. F., Linnemann, K. L., Bolz, S. N., Kaiser, F., Salentin, S., Haupt, V. J., & Schroeder, M. (2021). PLIP 2021: expanding the scope of the protein–ligand interaction profiler to DNA and RNA. Nucl. aci. res., 49(W1), W530–W534. 10.1093/nar/gkab294

59. Abraham, M. J., Murtola, T., Schulz, R., Páll, S., Smith, J. C., Hess, B., & Lindahl, E. (2015). GROMACS: High performance molecular simulations through multi-level parallelism from laptops to supercomputers. SoftwareX, 1, 19–25. 10.1016/j.softx.2015.06.001

60. Zoete, V., Cuendet, M. A., Grosdidier, A., & Michielin, O. (2011). SwissParam: a fast force field generation tool for small organic molecules. J. comp. chem., 32(11), 2359–2368. 10.1002/jcc.21816

61. Vanommeslaeghe, K., Hatcher, E., Acharya, C., Kundu, S., Zhong, S., Shim, J., Darian, E., Guvench, O., Lopes, P., Vorobyov, I., & MacKerell, A. D., Jr. (2010). CHARMM general force field: A force field for drug□like molecules compatible with the CHARMM all□atom additive biological force fields. J. comp. chem., 31(4), 671–690. 10.1002/jcc.21367

62. Zaki, A. A., Ashour, A., Elhady, S. S., Darwish, K. M., & Al-Karmalawy, A. A. (2022). Calendulaglycoside A showing potential activity against SARS-CoV-2 main protease: Molecular docking, molecular dynamics, and SAR studies. J. trad. comp. med., 12(1), 16–34. 10.1016/j.jtcme.2021.05.001

63. Daina, A., Michielin, O., & Zoete, V. (2017). SwissADME: a free web tool to evaluate pharmacokinetics, drug-likeness and medicinal chemistry friendliness of small molecules. Sci. rep., 7(1), 42717. 10.1038/srep42717

