## Supplementary material for "Design, Synthesis, Biological Evaluation and Molecular Docking of New Acid-Functionalized Carbazole Derivatives as Potential Antibiotic Agents": Suplementary

**Contents**

**Page No.**

1. **General Information	2**
2. **Figure S1. 1H NMR Spectra of 8	2**
3. **Figure S2. 13C NMR Spectra of 8	3**
4. **Figure S3. 13C DEPT Spectra of 8	3**
5. **Figure S4. 1H NMR Spectra of 4	4**
6. **Figure S5. 13C NMR Spectra of 4	4**
7. **Figure S6. 13C DEPT Spectra of 4	5**
8. **Figure S7. 1H NMR Spectra of 1	5**
9. **Figure S8. 13C NMR Spectra of 1	6**
10. **Figure S9. 1H NMR Spectra of 2	6**
11. **Figure S9. 1H NMR Spectra of 2	7**
12. **Figure S10. 13C NMR Spectra of 2	7**
13. **Figure S11. 1H NMR Spectra of 3	8**
14. **Figure S12. 13C NMR Spectra of 3	8**
15. **Figure S13. 13C DEPT Spectra of 3	9**
1. **General Information**

All the chemicals of reagent grade were purchased from several commercial sources and used without further purification. All reactions were performed under air and thin layer chromatography (TLC) was performed on Merck precoated TLC (silica gel 60 F_254_) plates. Melting points (^o^C) of the synthesised compounds were recorded in in open capillary tubes on a Digital melting point apparatus (Labard Scientific) and are uncorrected. The IR spectra were recorded in KBr discs on Schimadzu FTIR-8500. The ^1^H NMR (300 MHz) and ^13^C NMR (75 MHz) spectra were recorded on a Bruker Advance 300 spectrometer using CDCl_3_ or DMSO-d_6_ as the solvent. Data are reported as follows: chemical shift in ppm (δ), multiplicity (s = singlet, d = doublet, t = triplet, q = quartet, and m = multiplet), coupling constant (Hz). High-resolution mass spectra (HRMS) data were recorded on Qtof Micro YA263.

**1. ^1^H and ^13^C NMR Spectra of (*E*)-4-(2-(4-methyl-2-oxocyclohexylidene)hydrazinyl)benzoic acid (8):**

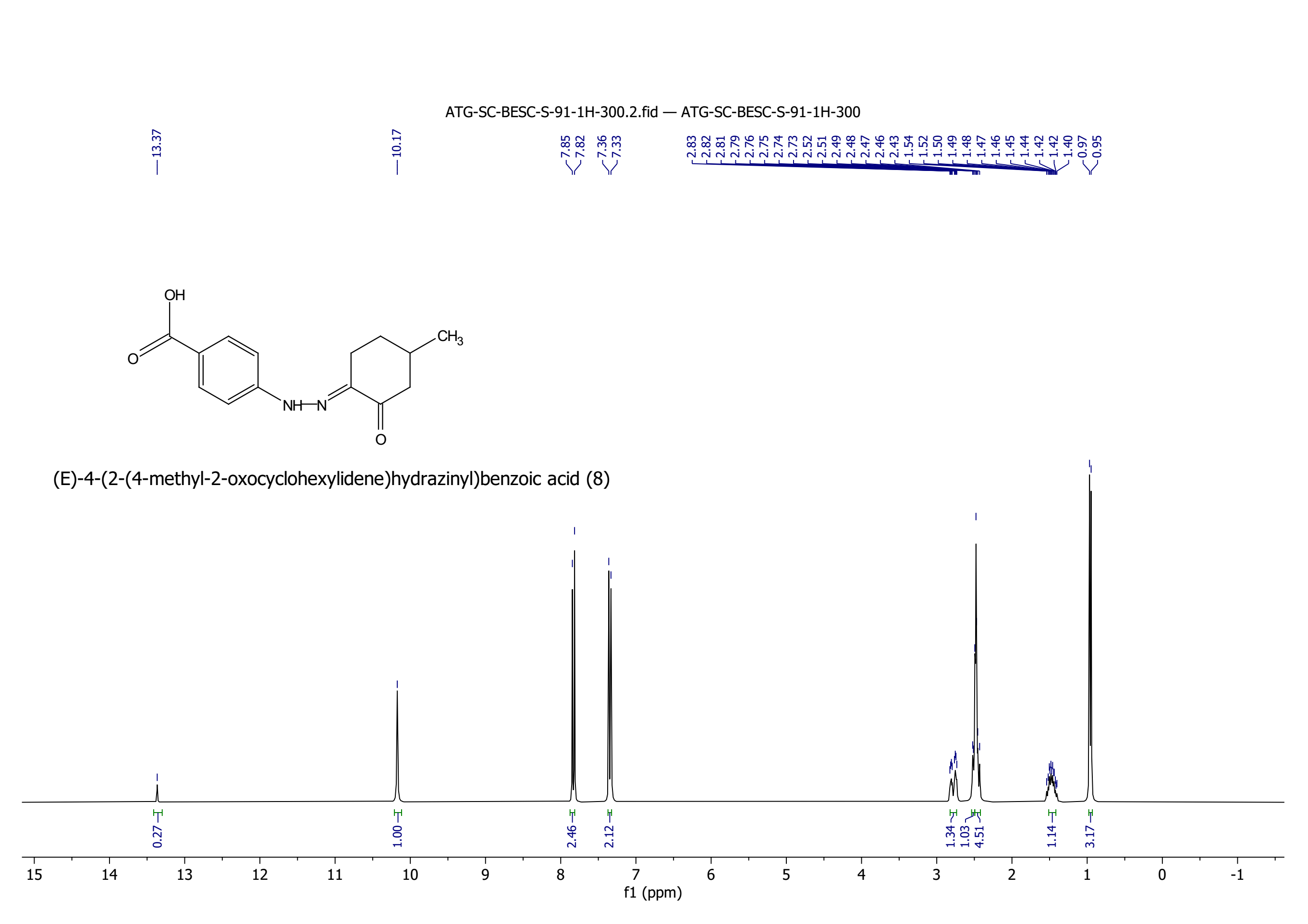

**Figure S1. ^1^H NMR Spectra of 8**

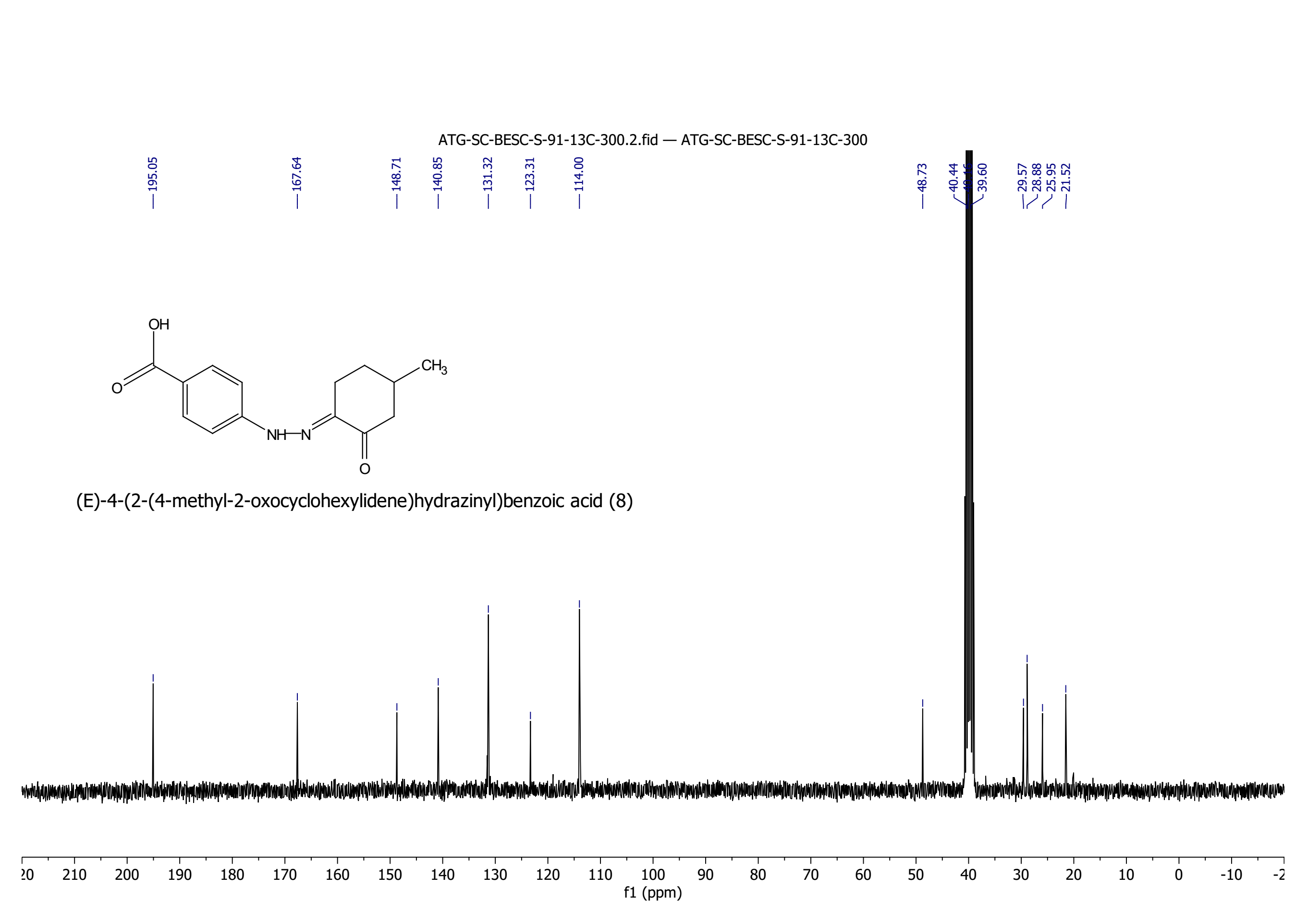

**Figure S2. ^13^C NMR Spectra of 8**

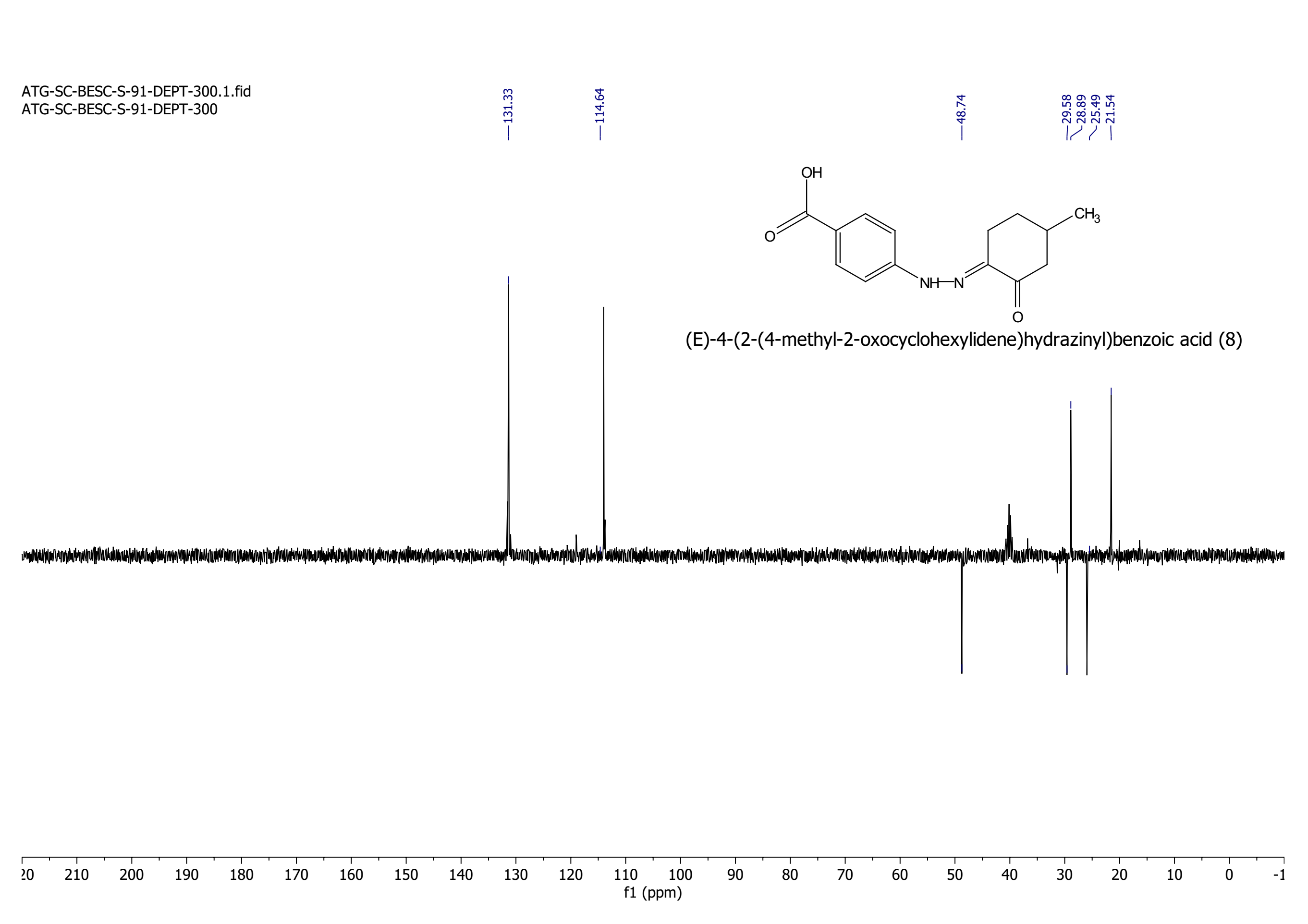

**Figure S3. ^13^C DEPT Spectra of 8**

**2. ^1^H and ^13^C NMR Spectra of 3-methyl-1-oxo-2,3,4,9-tetrahydro-1*H*-carbazole-6-carboxylic acid (4):**

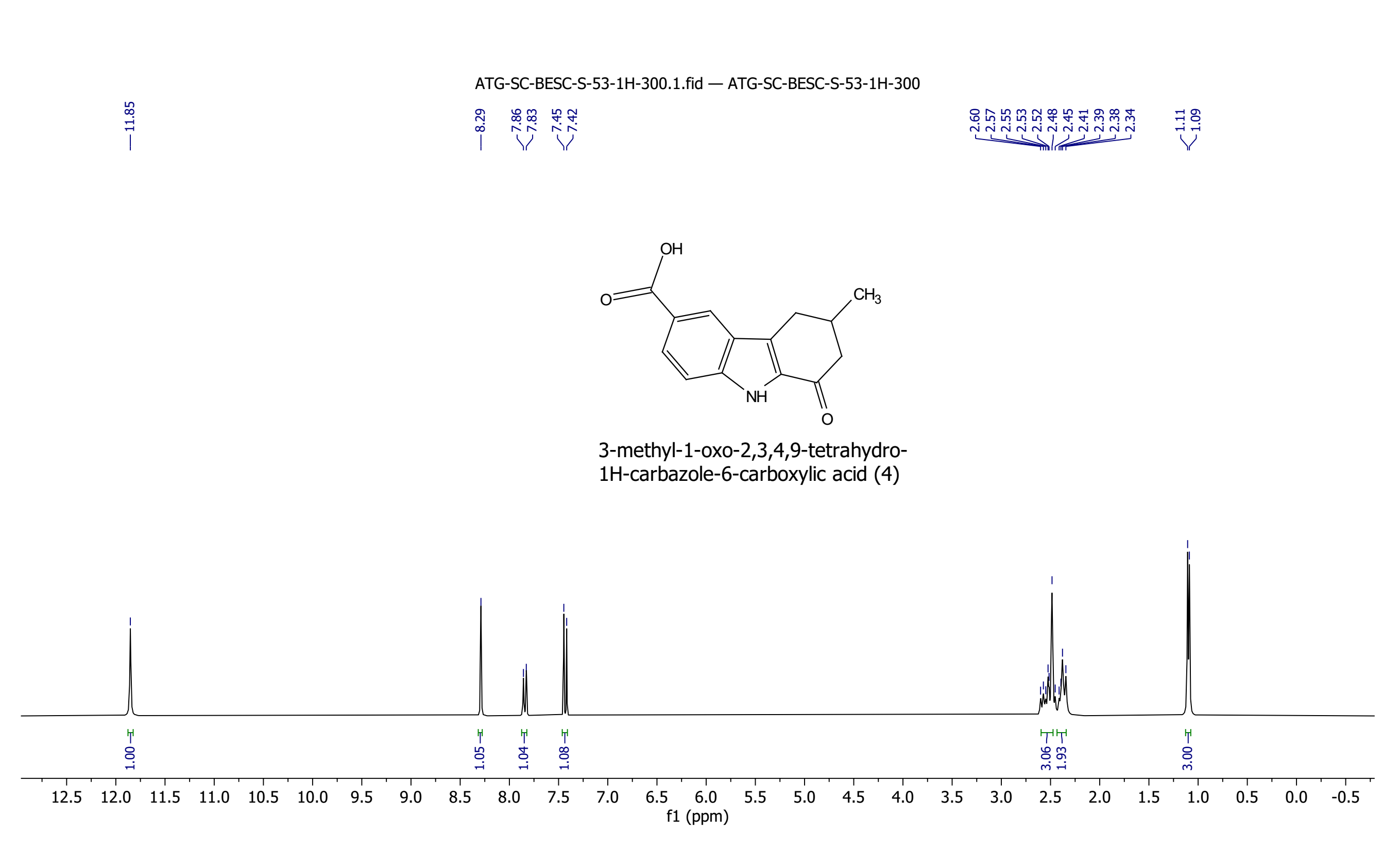

**Figure S4. ^1^H NMR Spectra of 4**

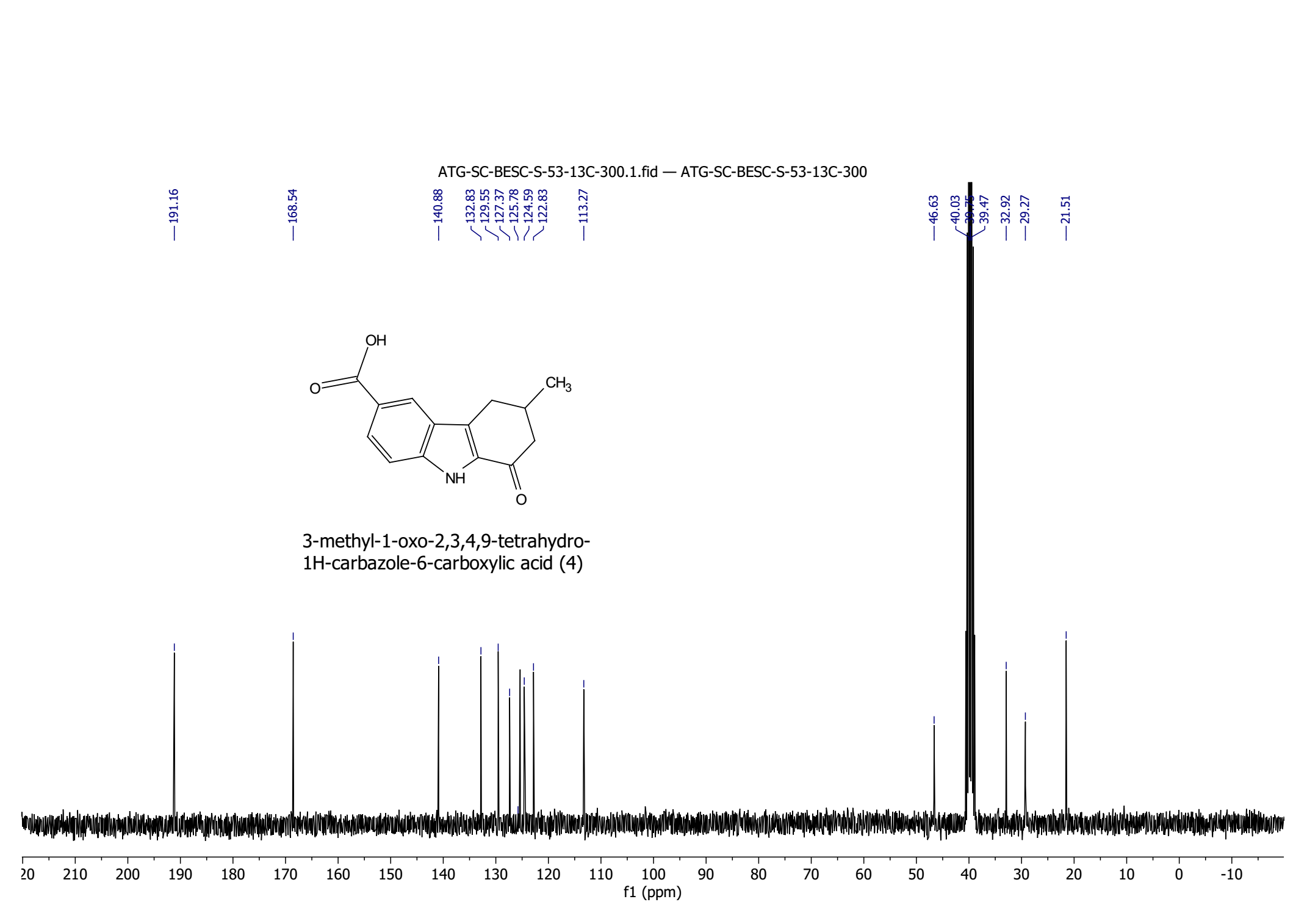

**Figure S5. ^13^C NMR Spectra of 4**

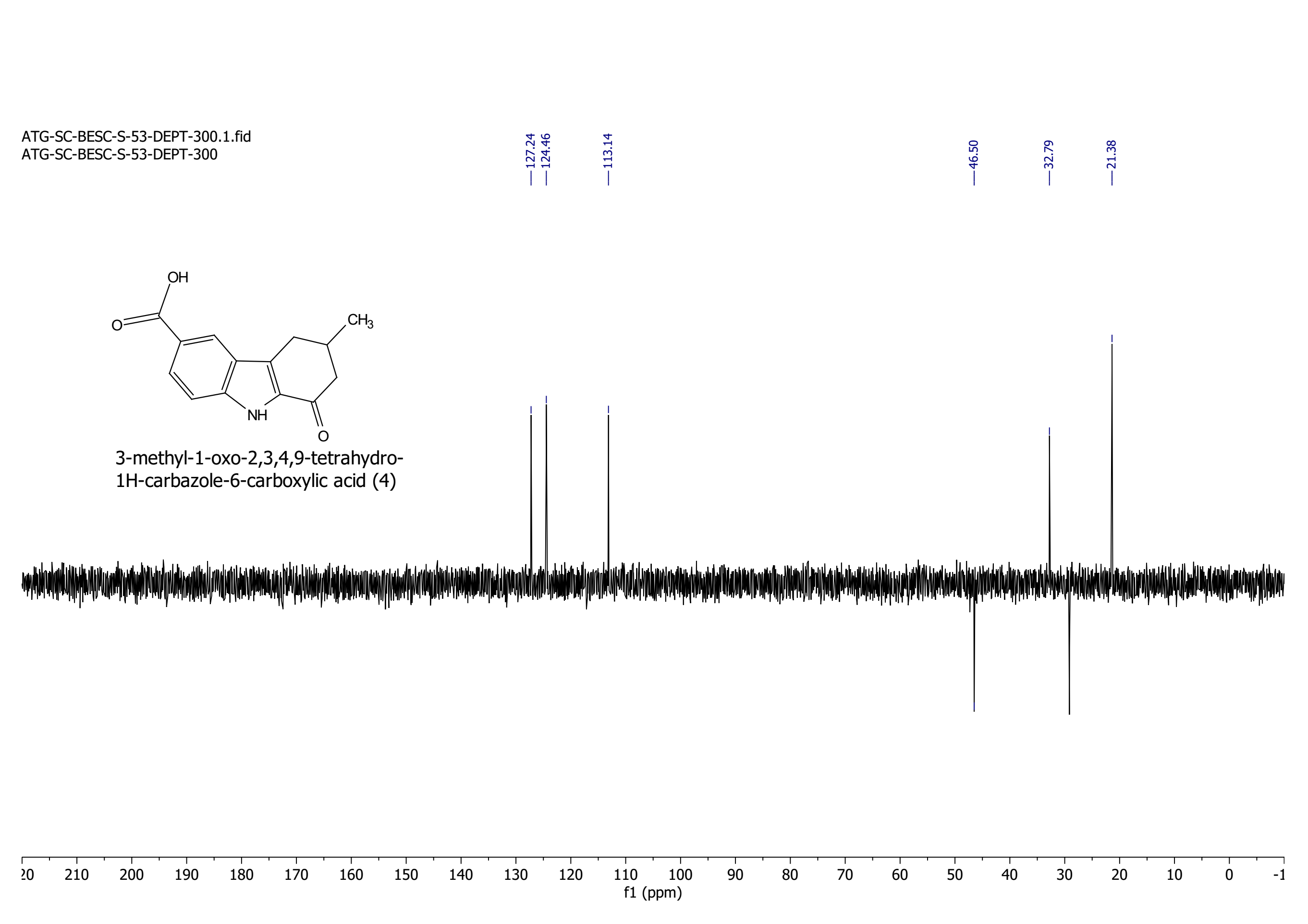

**Figure S6. ^13^C DEPT Spectra of 4**

**3. ^1^H and ^13^C NMR Spectra of (*E*)-3-methyl-1-(2-tosylhydrazono)-2,3,4,9-tetrahydro-1*H*-carbazole-6-carboxylic acid (1):**

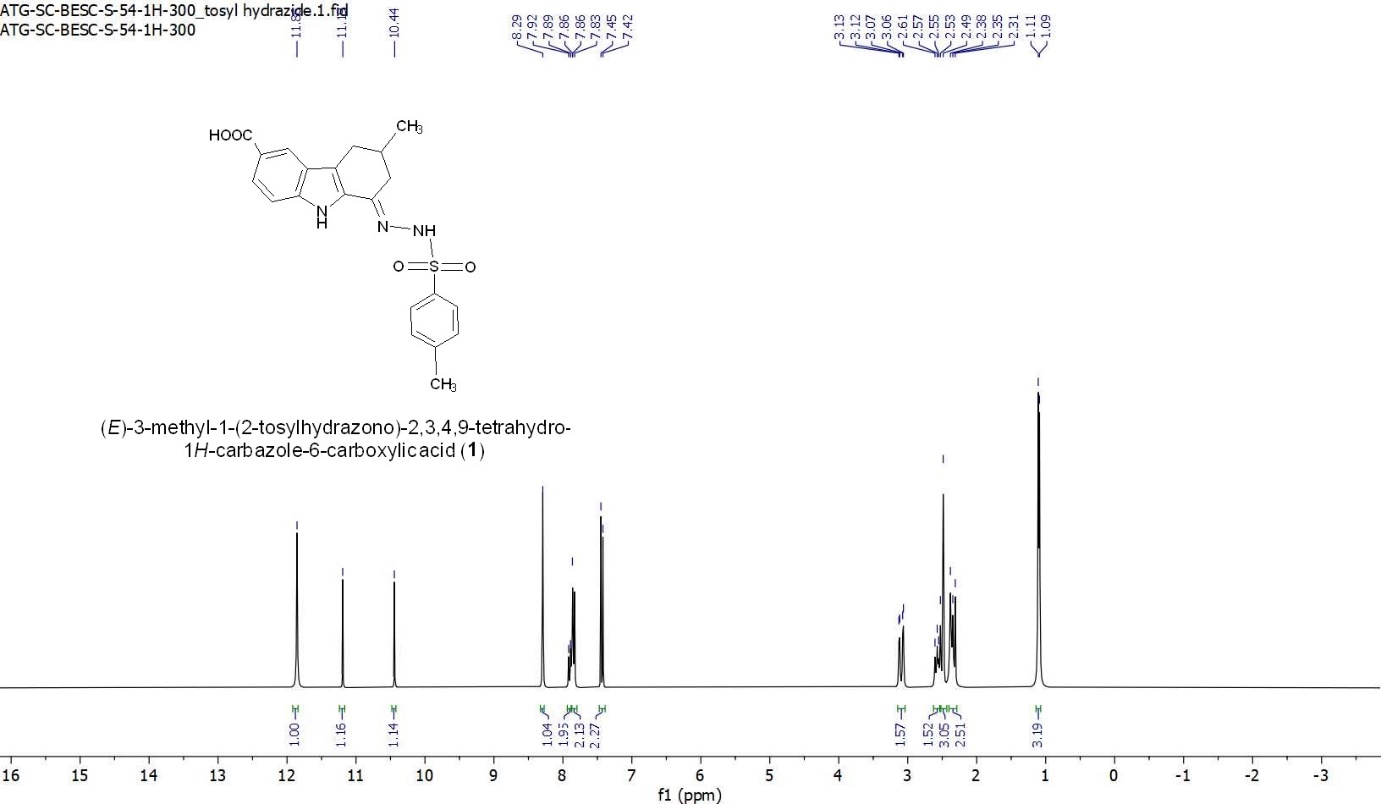

**Figure S7. ^1^H NMR Spectra of 1**

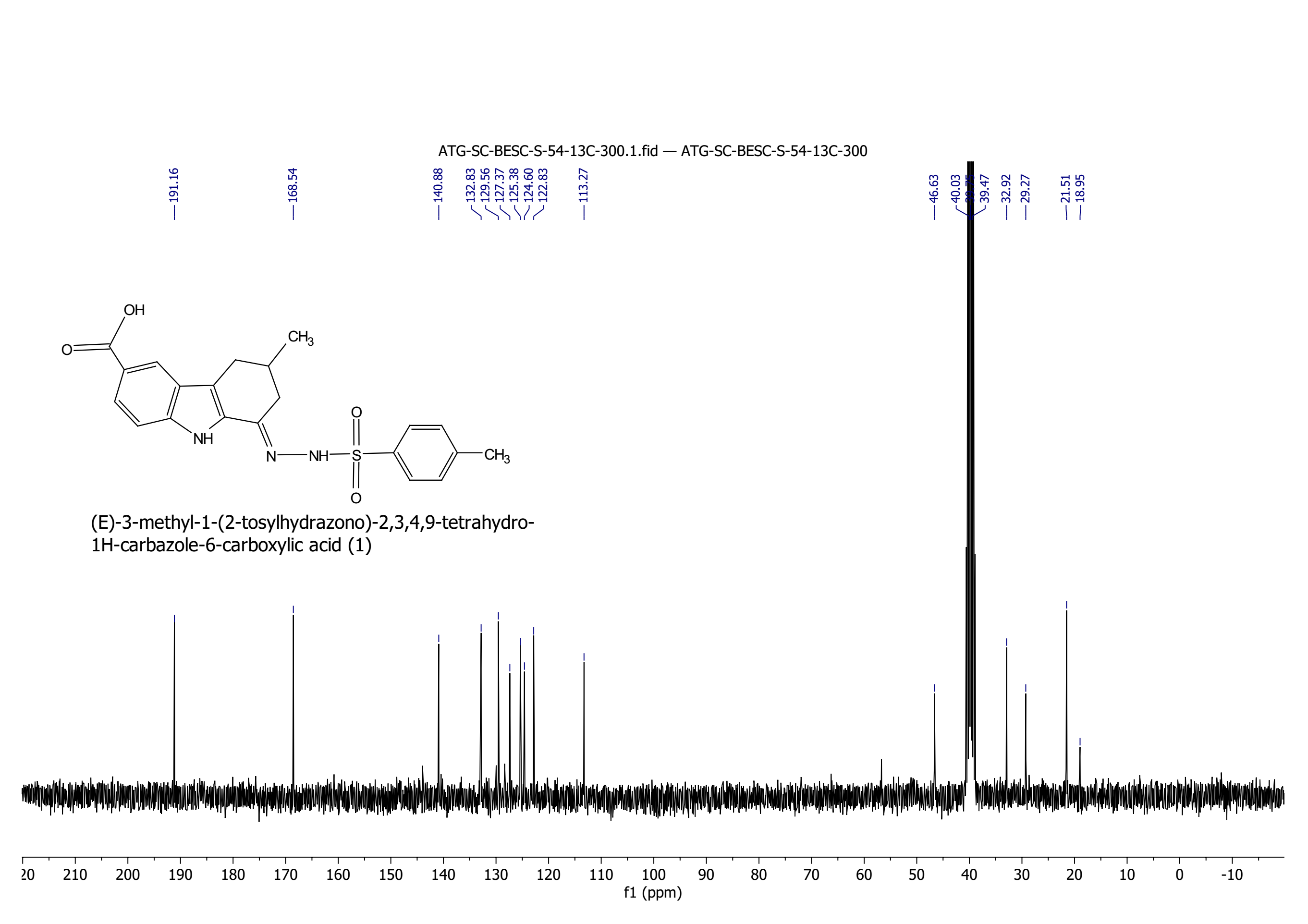

**Figure S8. ^13^C NMR Spectra of 1**

**4. ^1^H and ^13^C NMR Spectra of 3-methyl-1,4-dioxo-4,9-dihydro-1*H*-carbazole-6-carboxylic acid (2):**

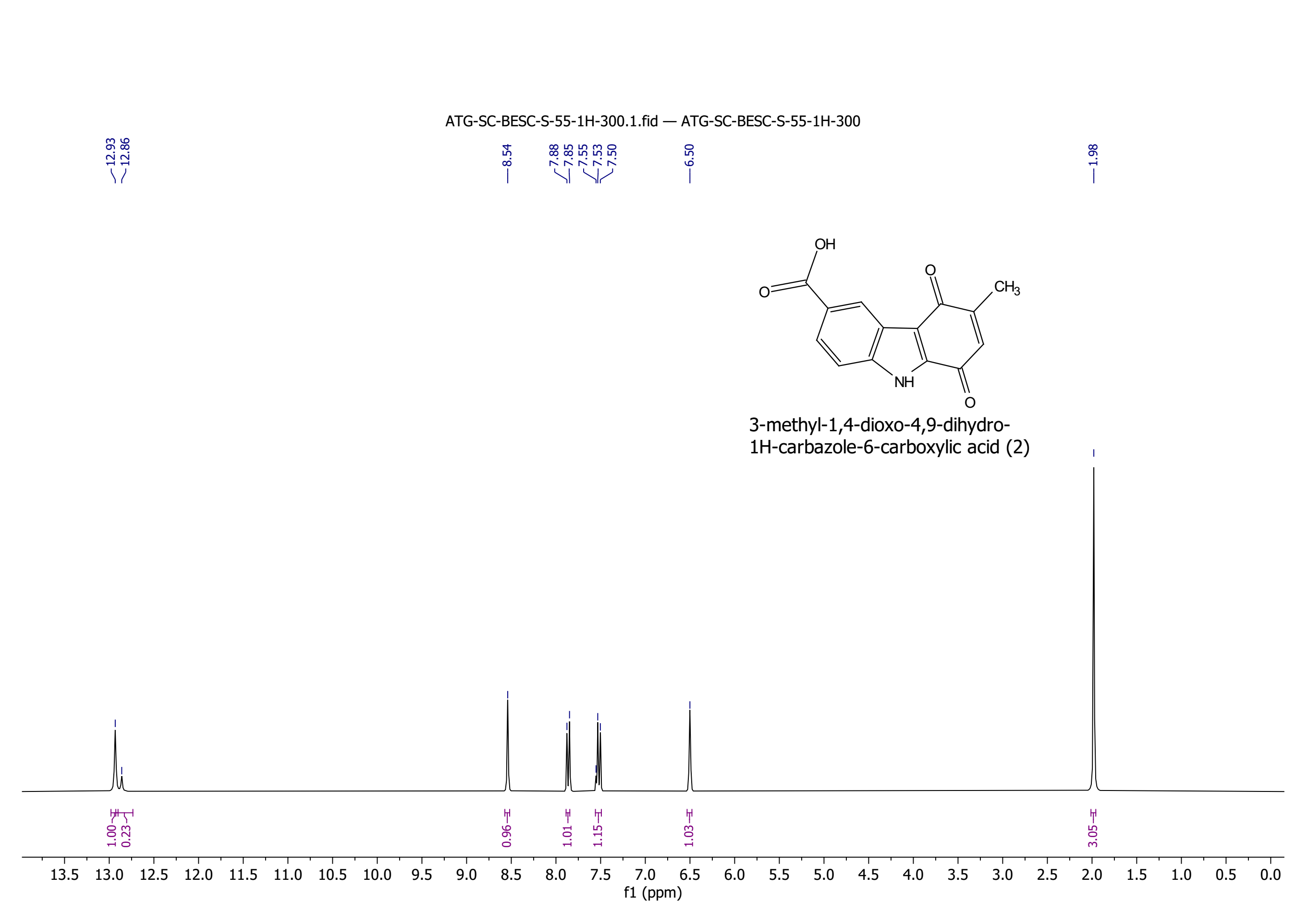

**Figure S9. ^1^H NMR Spectra of 2**

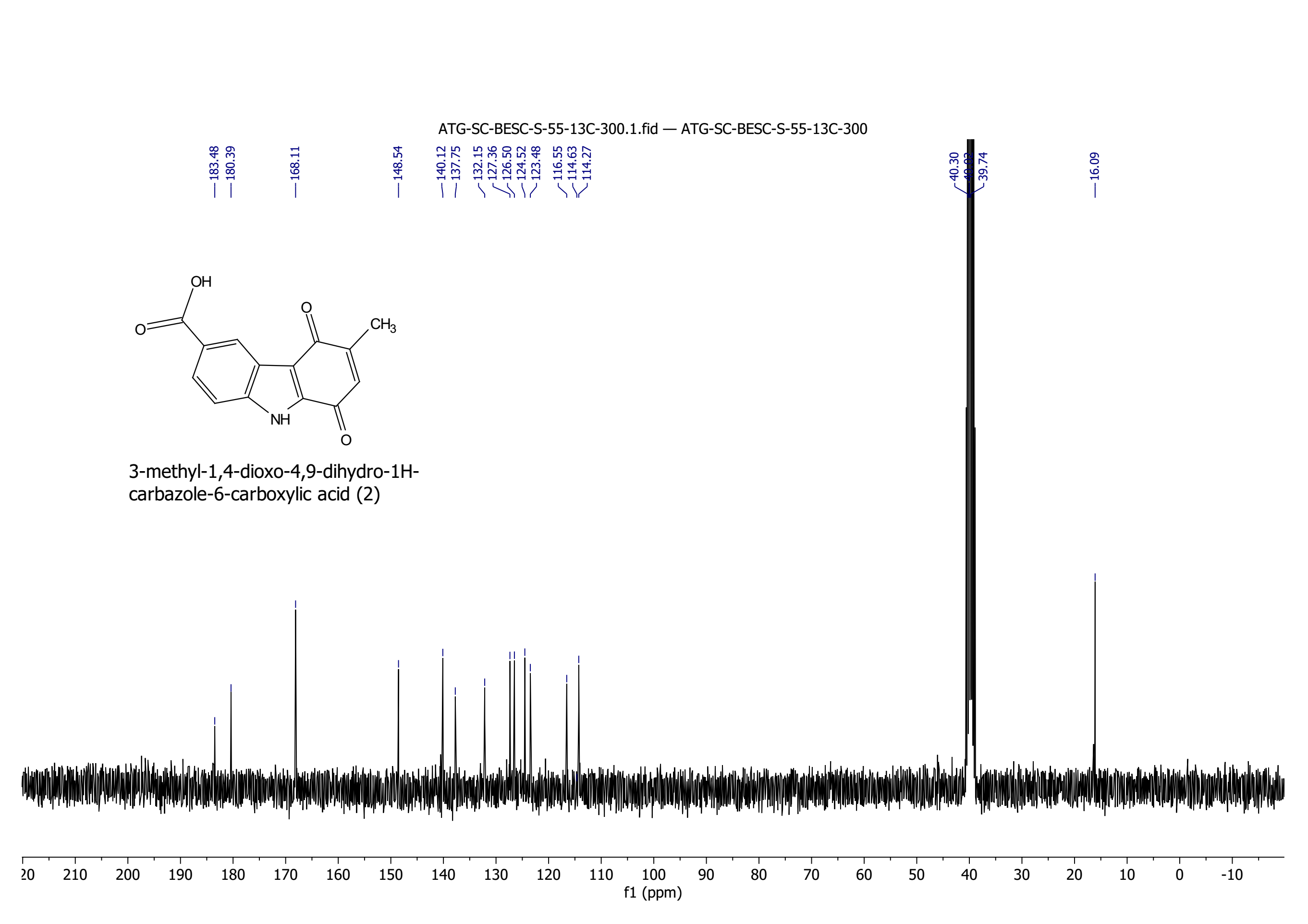

**Figure S10. ^13^C NMR Spectra of 2**

**5. ^1^H and ^13^C NMR Spectra of 6-methyl-9*H*-carbazole-3-carboxylic acid (3):**

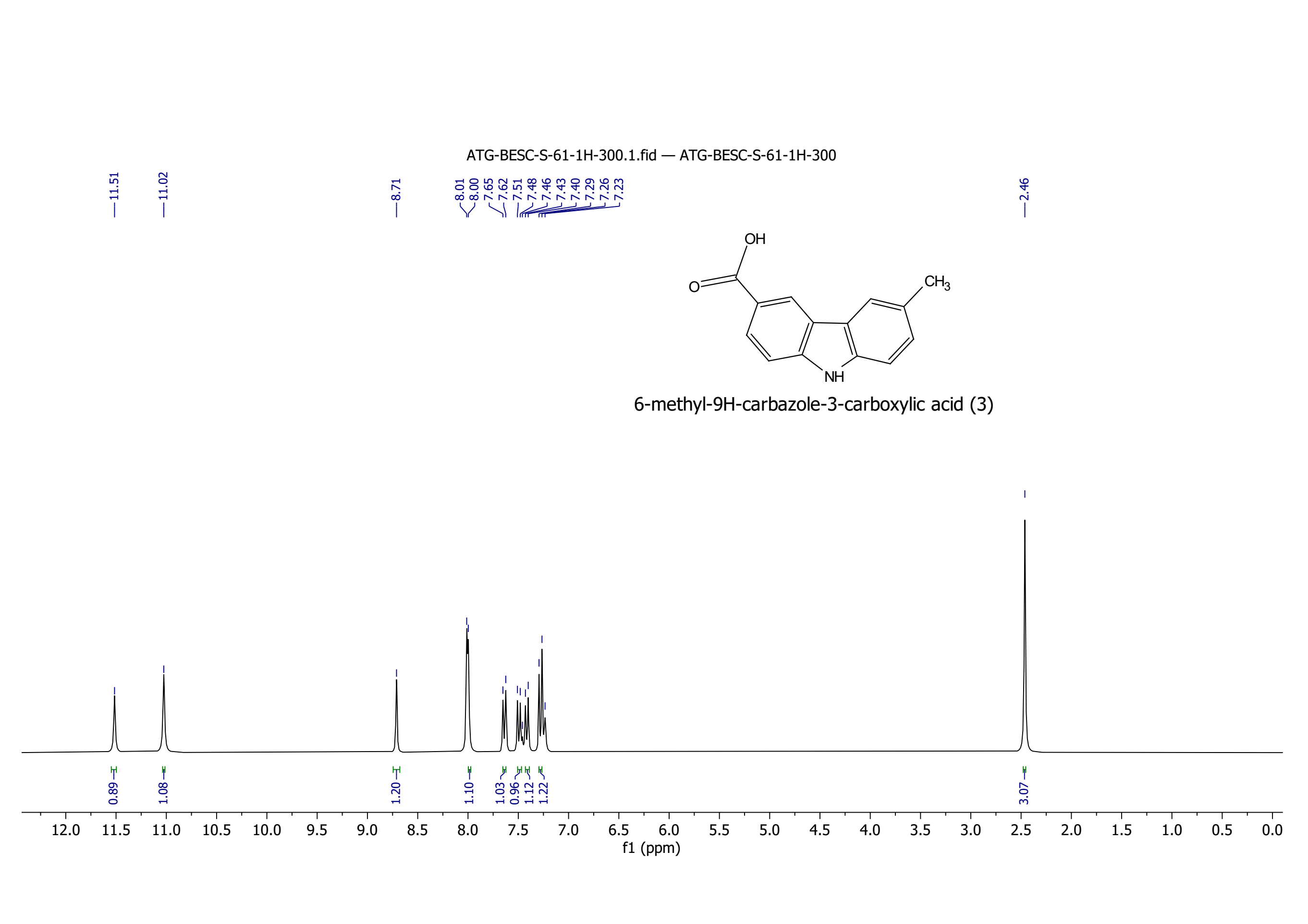

**Figure S11. ^1^H NMR Spectra of 3**

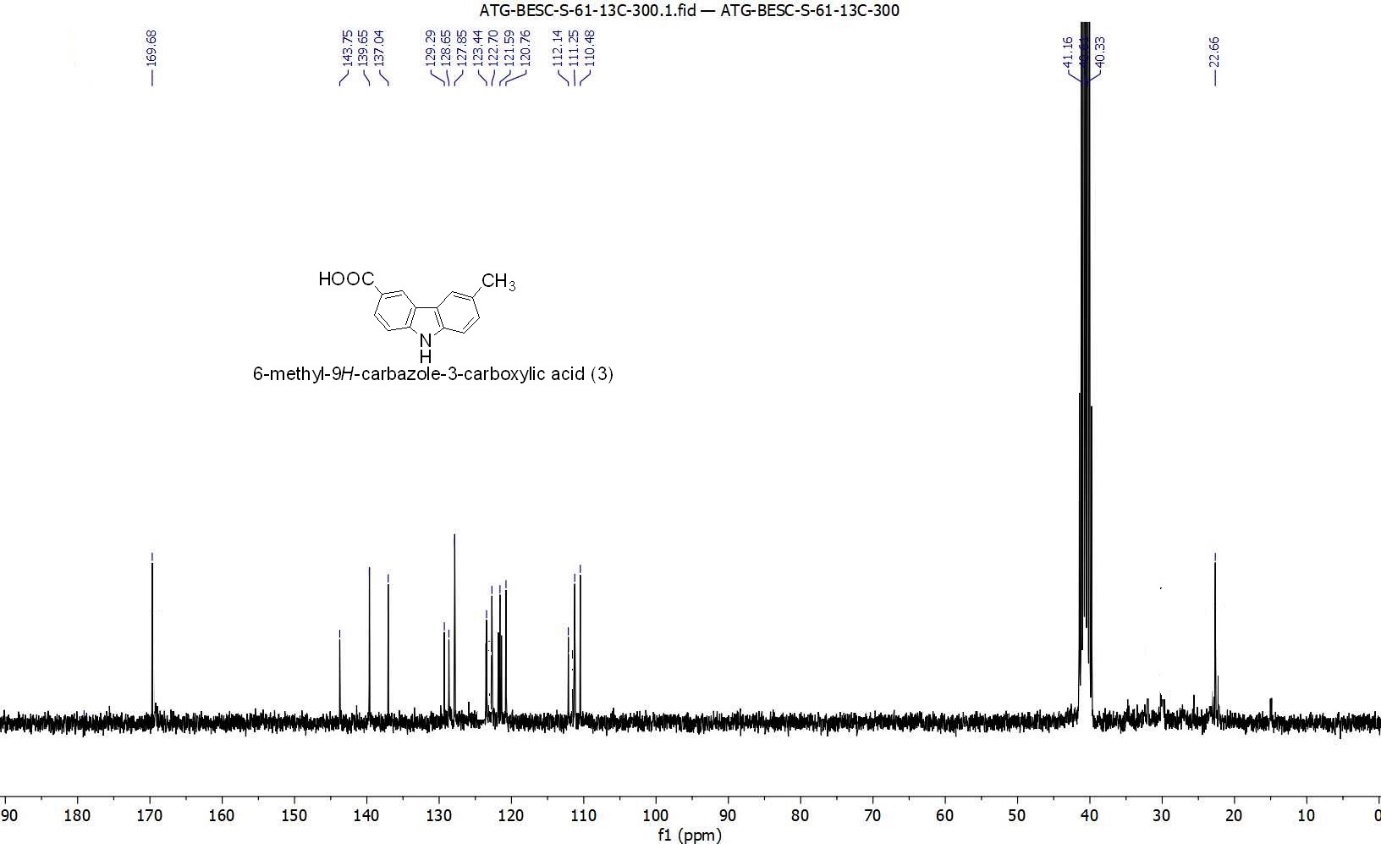

**Figure S12. ^13^C NMR Spectra of 3**

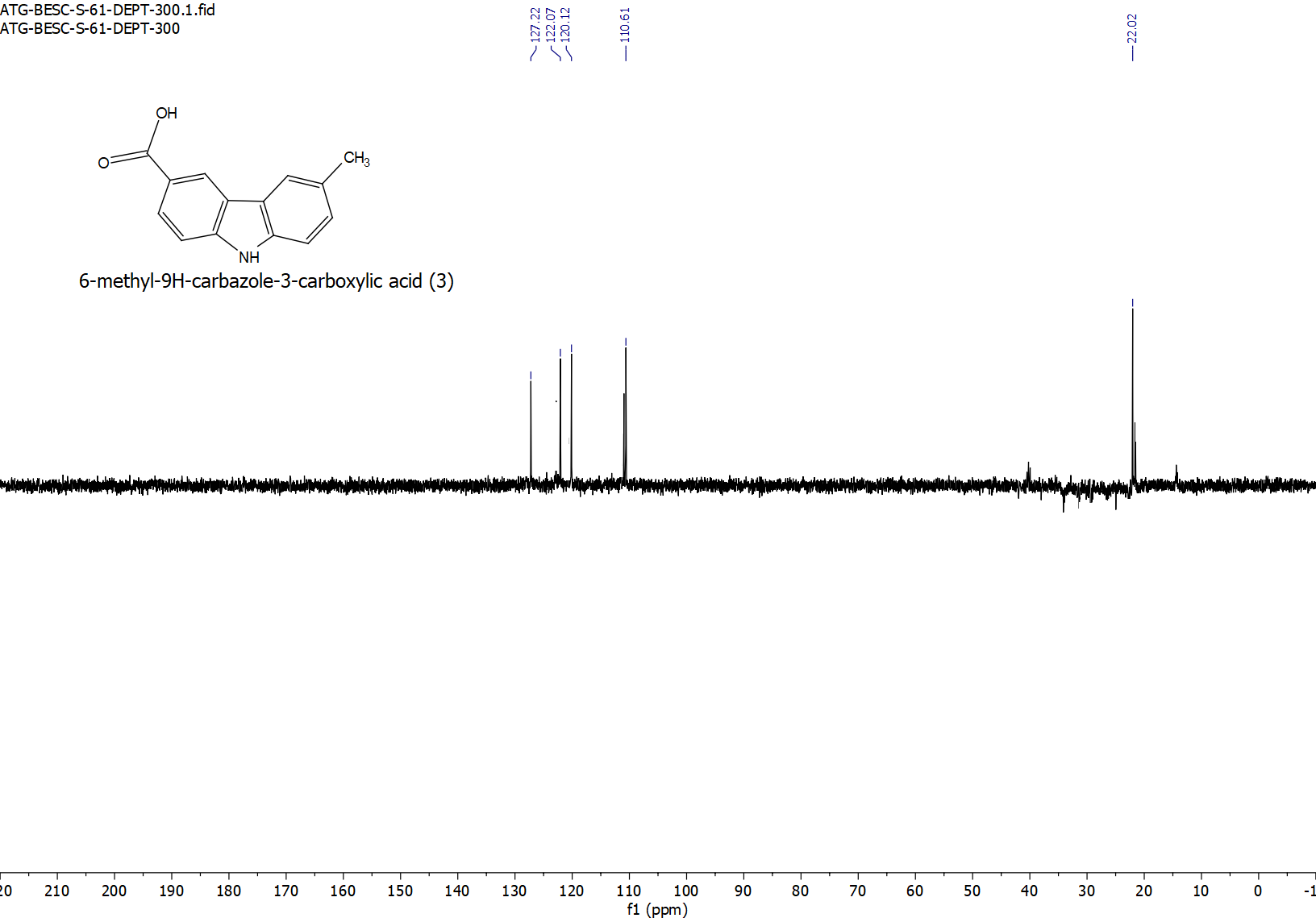

**Figure S13. ^13^C DEPT Spectra of 3**
